# BMP9 regulates the endothelial secretome to drive pulmonary hypertension

**DOI:** 10.1101/2025.08.29.673113

**Authors:** Ying Zhong, Peiran Yang, Luca Troncone, Oleg V. Kovalenko, Eva M. Fast, Shreyas Rajesh, Sydney Lavoie, Kathleen E. Tumelty, Elizabeth Shin, Taylor A. Covington, Ana Zeghibe, Marc H. Wadsworth, Stephen M. Christensen, Robin Nathans, Susan A. Benard, Xianchun Tang, Shakey A. Quazi, Lillian R. Worst, Megan E. McNeil, Stephanie S. Kim, Geoffrey D. Bocobo, Robert Szulcek, Harm J. Bogaard, Kevin M. Hart, Erik M. Martinez-Hackert, Stephen P. Berasi, Christine Huard, Paul B. Yu

## Abstract

BMP9, a pleiotropic growth factor cytokine that regulates endothelial function, is implicated in the pathogenesis of pulmonary arterial hypertension (PAH). Loss-of-function mutations in *GDF2* are found in heritable PAH, suggesting its function as an endothelial quiescence factor, while agonizing or antagonizing BMP9 signaling are both reported to ameliorate experimental pulmonary hypertension (PH). This study sought to resolve the contribution of BMP9 to pulmonary vascular disease and its status as a potential therapeutic target. The function of BMP9 in experimental PH was interrogated using recombinant BMP9, BMP9/BMP10 ligand trap ALK1-Fc, two anti-BMP9 neutralizing antibodies, and the activin/GDF/BMP ligand trap ACTRIIA-Fc (a.k.a., sotatercept). Disulfide-linked, prodomain complexed BMP9 was not protective in SUGEN-hypoxia or monocrotaline-induced PH models, in contrast to previous studies using incompletely disulfide-linked BMP9. In comparison, selective and non-selective BMP9 antagonism exerted prophylactic and therapeutic effects across PH models. Anti-BMP9 and ACTRIIA-Fc had comparable impact on hemodynamics, RV hypertrophy, and vascular remodeling, while single nucleus RNA-Seq revealed similar inhibition of SMAD1/5 and SMAD2/3 transcriptional activity, and highly overlapping DEGs, particularly in the endothelial compartment (r=0.83, p=2.54e-43, Spearman), suggesting overlap of mechanism in targeting BMP9. A multi-omic approach using lung tissues from human PAH, experimental models of pulmonary hypertension, and transcriptomic analysis of pulmonary microvascular endothelial cells from PAH patients revealed that BMP9 is critical for regulating several endothelial gene products that are overexpressed in human and experimental disease and implicated in disease pathogenesis including *CXCL12, PDGF-BB, EDN1, COL18A1*, and *IGFBP4*, and are inhibited by administering anti-BMP9 neutralizing antibodies or ligand traps. Co-culture studies revealed paracrine effects of BMP9-stimulated PMVEC on pulmonary arterial smooth muscle cell (PASMC) phenotypic plasticity, which could be attributed in large part to endothelial-derived CXCL12. In summary, endothelial BMP9 signaling is a key coordinator of vasoactive endothelial gene products that modulate PASMC phenotype and appears to be a shared target of anti-BMP9 and ACTRIIA-Fc. Selective targeting of endothelial BMP9 angiogenic signaling represents a potential therapeutic strategy for human PAH.

**One sentence summary:** Inhbiition of BMP9 signaling via different selective and non-selective strategies, including ACTRIIA-Fc/sotatercept, attenuates endothelial expression of vasoactive secreted factors that drive pulmonary vascular remodeling.

## Introduction

Pulmonary arterial hypertension (PAH), defined hemodynamically by a mean pulmonary arterial pressure of >20 mmHg, a pulmonary vascular resistance of > 2 Woods Units, and a pulmonary capillary wedge pressure of ≤15 mmHg,^1^ is a highly morbid disorder of elevated pulmonary vascular resistance marked by progressive obliteration of the pulmonary arteriolar circulation. Survival with currently available therapies remains slightly better than 50% at 5 years, with mortality resulting from progression of PAH and subsequent right heart failure.^2,3^ Currently approved vasodilator therapies improve functional status and composite survival endpoints, but are limited by systemic effects and tachyphylaxis, and are not designed to target underlying mechanisms of pulmonary vascular remodeling and vessel loss. The advent of sotatercept, a.k.a., the activin type II receptor extracellular domain Fc fusion protein (ACTRIIA-Fc), introduced the concept of an activin/GDF/BMP-targeted therapy for the treatment of PAH,^4^ demonstrating improvements in hemodynamics, function, and survival as add-on therapy,^5–8^ suggesting modulation of these signaling pathways has potential for modifying the disease process.

The pathogenetic role of BMP/TGFβ signaling in PAH is supported by loss-of-function (LOF) mutations associated with heritable PAH are found in genes encoding the BMP type 2 receptor *BMPR2*, the BMP type 1 receptor *ACVRL1/*ALK1, co-receptor *ENG/*endoglin, ligand *GDF2*/BMP9, and downstream effector *SMAD9*—describing the endothelial BMP9-BMPR2-ALK1-ENG signal transduction complex. Moreover, the role of these LOF loci in PAH are consistent with the critical roles attributed to BMP9 in regulating vascular development, endothelial homeostasis and barrier function^9–12^. Diminished circulating BMP9 is found in portopulmonary hypertension^13,14^, also suggesting a key role of BMP9 signaling in pulmonary vascular homeostasis. Mechanistic studies on the role of BMP9 in experimental pulmonary hypertension (PH) have been less consistent. Treatment with recombinant BMP9 attenuates PH and pulmonary vascular remodeling in several animal models,^15^ whereas BMP9-deficient mice are partially protected from experimental PH, and administration of BMP9/BMP10 ligand trap ALK1-Fc or neutralizing antibody to BMP9 are also protective.^16^ These divergent findings have suggested that BMP9 signaling may have context-sensitive effects in pulmonary vascular biology. Consistent with context-sensitive effects, BMP9 exerts anti-proliferative effects in wild-type pulmonary vascular endothelial cells that are lost in BMPR2 haploinsufficient cells, and exerts pro-proliferative effects in BMPR2 homozygous deficient cells.^17^ Similarly, we have found that BMP9 exerts anti-proliferative effects in pulmonary microvascular endothelial cells (PMVEC) obtained from non-PAH donor lungs, and pro-proliferative effects in PMVEC obtained from explanted PAH lungs.^4^

To understand the contribution of BMP9 vs. related ligands in pulmonary vascular disease, we evaluated the effects of two anti-BMP9 neutralizing antibodies, BMP9/10 ligand trap ALK1-Fc, and activin/GDF/BMP9/BMP10 ligand trap ACTRIIA-Fc, versus a well-characterized recombinant BMP9 pro-complex protein in several models of experimental PH. Whole tissue (bulk) and single nucleus (sn) RNA-seq studies using treated or untreated experimental PH lung tissues, and in PMVEC exposed to BMP9 were performed to reveal potential mechanisms of action. To capture potential paracrine effects of endothelial BMP9 signaling on apposed pulmonary smooth muscle cells (PASMC), signaling and functional studies were performed in PMVEC-PASMC co-culture models. These studies reveal overlapping mechanisms of action of anti-BMP9, ALK1-Fc, and ACTRIIA-Fc in modifying endothelial function, suggesting that each may elicit effects from neutralizing BMP9. BMP9 is a pleiotropic growth factor cytokine that regulates a large number of secreted vasoactive gene products in the pulmonary microvascular endothelium, and may serve as a master regulator of angiogenic signaling to drive pulmonary vascular remodeling in PH.

## Results

### Inhibition of BMP9/BMP10 attenuates experimental PH before or after pulmonary vascular injury

A previous report showed BMP9/10 ligand trap ALK1-Fc to be protective when administered to monocrotaline (MCT-) or SU5416 and hypoxia (SU-Hx-) exposed rats after the development of experimental PH.^16^ We sought to test if the effects of BMP9/BMP10 inhibition might depend on whether ALK1-Fc is administered before or after the development of PH. Consistent with prior observations, administering ALK1-Fc (10 mg/kg s.c. twice weekly) for 3 weeks after the establishment of severe PH due to SU5416 (20 mg/kg s.c.) and hypoxia (FIO2=0.1) significantly reduced right ventricular systolic pressure (RVSP) and right ventricular hypertrophy (RVH) based on Fulton’s index (RV/LV+S, “post-treatment”, **Fig. 1A-C**). Administration of ALK1-Fc for 1 week prior to exposure to SU-Hx did not exacerbate experimental PH but elicited a trend towards reduced RVSP (“pre-treatment”, **Fig 1A-C**). Next, we tested whether or not suppression of BMP9/BMP10 signaling via ALK1-Fc might synergize with hypoxia to exacerbate PH and vascular remodeling. Administration of ALK1-Fc coinciding with exposure to hypoxia did not exacerbate, but rather attenuated PH and RVH due to hypoxia, resulting in essentially normal RVSP and Fulton’s indices (**Fig. 1D-F**). Histomorphometry revealed that treatment with ALK1-Fc before or after exposure SU-Hx significantly improved pulmonary small arteriolar muscularization based on smooth muscle actin (SMA) staining of small (<50 μm) vessels as compared to SU-Hx treatment alone (**Fig. 1G-H**). Moreover, the combination of ALK1-Fc with hypoxia resulted in less muscularization than SU-Hx-induced remodeling and was statistically not different than normoxia. These hemodynamic and histologic results demonstrate inhibiting BMP9/BMP10 signaling improves experimental PH whether it is administered before, concurrently, or after the development of experimental PH. Despite varying the timing of BMP9/BMP10 in relation to pulmonary vascular injury, these experiments revealed a pathogenic rather than protective role of BMP9/BMP10 signaling in all contexts tested.

**Figure 1.**
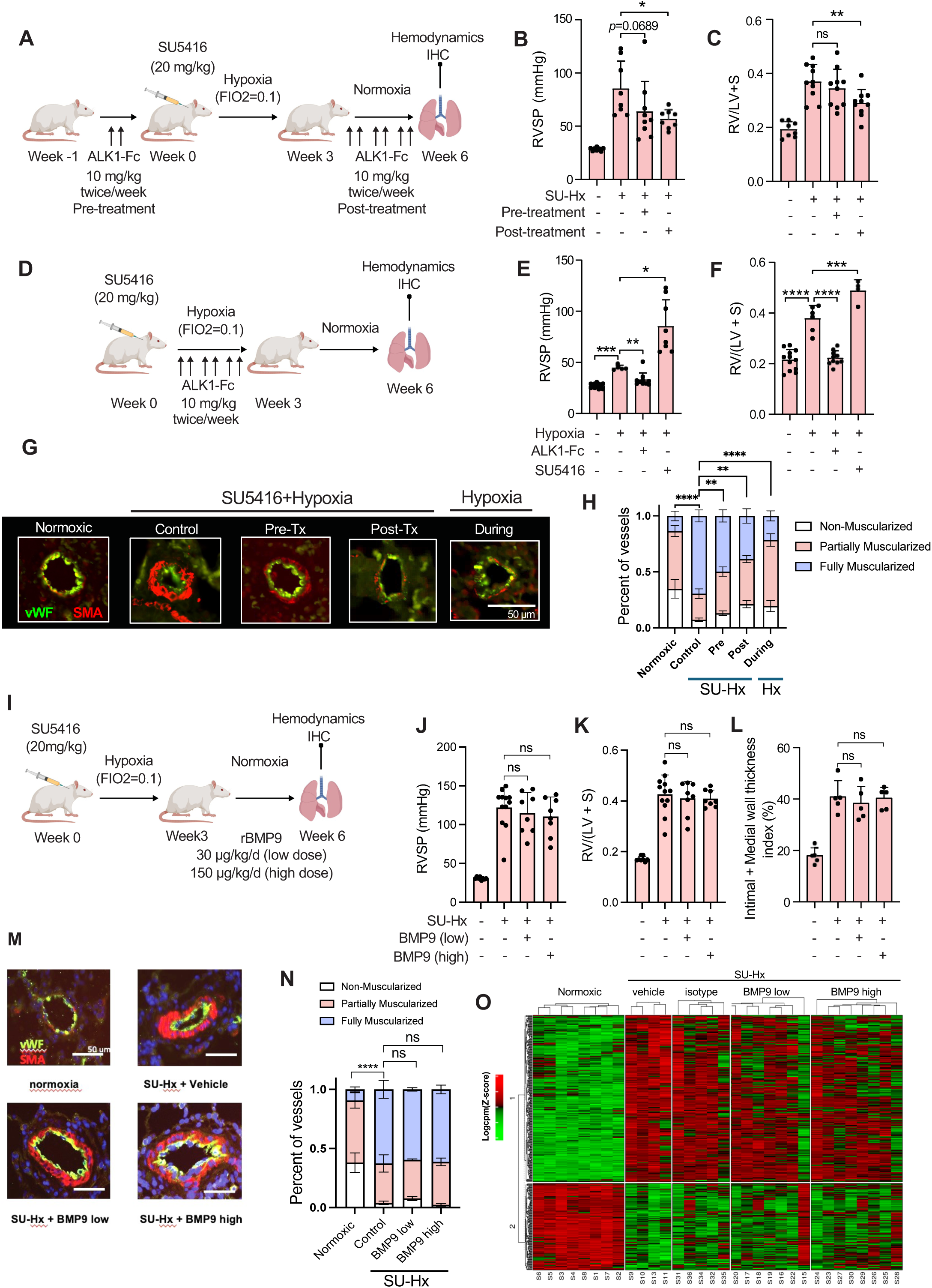
The impact of BMP9/BMP10 ligand trap ALK1-Fc at varying intervals in relation to SU-Hx or hypoxia in SD rats. **(A)** Administering ALK1-Fc (6 doses) after the development of PH 3 weeks following exposure to SU-Hx elicited improvements in **(B)** RVSP and **(C)** Fulton’s index as compared to vehicle-treated rats, whereas administering ALK1-Fc (2 doses) before exposure to SU-Hx elicited a trend towards improved RVSP. (*n* = 8 to 10 per group, mean ± SD, *P<0.05, **P<0.01, One-way ANOVA with Sidak’s test. **(D)** Administering ALK1-Fc (6 doses) coinciding with exposure to hypoxia elicited improvements in **(E)** RVSP and **(F)** Fulton’s index as compared to hypoxia-only exposed rats. (*n* = 6 – 12 per group, mean ± SD, *P<0.05, **P<0.01, ***P<0.001, ****P<0.0001, One-way ANOVA with Dunnett’s test (RVSP) and Sidak’s Test (Fulton’s index). **(G)** Immunofluorescence images reveal muscularization of small (<50 μm) pulmonary arterioles in SU-Hx vs. normoxic rats, which was attenuated by ALK1-Fc administered before (Pre-Tx), after (Post-Tx), or during hypoxia. **(H)** The percentage of fully, partially and non-muscularized vessels in normoxic rats vs. SU-Hx rats pre- or post-treated with ALK1-Fc compared to vehicle control and Hx rats treated with ALK1-Fc (diameter, 10 to 50 µm). Values shown mean ± SD, \*\**P*<0.01, \*\*\*\**P*<0.0001, one-way ANOVA with Tukey’s test for the percentage of fully muscularized vessels). **(I)** Adult SD rats were subjected to SU-Hx and (SUGEN5416 20 mg/kg, sc and FiO2 = 0.10) for 3 weeks, followed by 3 weeks of treatment in normoxia with rBMP9 (30 µg/kg for low dose, 150 µg/kg for high dose, ip, daily) and the examined for **(J)** RVSP, **(K)** Fulton’s index and **(L)** intimal-medial thickness index of pulmonary arterioles < 50 µm (*n* = 5 to 11 per group, means ± SD, \**P* < 0.05, one-way ANOVA with Dunnett’s test. **(M)** Immunofluorescence images of muscularization of pulmonary arterioles in rats subjected to normoxia, SU-Hx, treated with vehicle, low or high dose rBMP9. **(N)** The percentage of fully, partially and non-muscularized vessels (diameter <50 µm, mean ± SD, two-way ANOVA with Dunnett’s test for percentage of fully muscularized vessels). **(O)** Heatmap showing the lung transcriptomes of Nx rats, SU-Hx rats and SU-Hx rats treated with vehicle, low dose BMP9, or high dose BMP9.

### Treatment with validated recombinant pro-complex BMP9 does not attenuate established experimental PH

Based on the rationale that HPAH, non-genetic PAH, and animal PH model lung tissues exhibit diminished *BMPR2* expression and BMP signaling,^18,19^ systemic treatment with exogenous recombinant BMP9 (rBMP9) was proposed, and found to attenuate experimental PH and pulmonary vascular remodeling in MCT-treated rats, mutant *Bmpr2* mice, and SU-Hx treated mice and rats, attenuating vessel loss in rats with severe obliterative PH due to SU-Hx,^15^ supporting the concept that BMP9 exerts a protective role in PH. To revisit this concept, we used rBMP9 expressed as a mature BMP9 homodimer in complex with its prodomain, or “pro-complex rBMP9”, to recapitulate the previous study and to represent the circulating form of BMP9 homodimer. In contrast to previous findings, administration of pro-complex rBMP9 (600 ng/day s.c., ∼3 µg/kg/d, as before^15^) for 2 weeks starting 2 weeks after treatment of rats with MCT did not elicit significant changes in RVSP, RVH or pulmonary arteriolar muscularization versus vehicle-treated controls (**Fig. S1A-F**), a result replicated in 3 independent experiments using 2 distinct batches of pro-complex rBMP9. Similarly, administration of pro-complex rBMP9 at higher doses (30 µg/kg/d, 150 µg/kg/d s.c., ∼10-fold and 50-fold previously used doses) for 3 weeks after establishment of SU-Hx PH in rats did not elicit significant decreases in RVSP, RVH, or pulmonary arteriolar muscularization versus vehicle controls (**Fig. 1I-N**). Similarly, treatment with pro-complex rBMP9 at either dose did not elicit changes in gene expression versus treatment with vehicle or isotype Ab in lung tissues from SU-Hx treated rats based on bulk RNA-Seq analysis (heatmaps, **Fig. 1O**).

Given differences in the impact of pro-complex rBMP9 in current versus previous studies, we sought to characterize biochemical differences in rBMP9 protein preparations that might account for discrepant results. New and previous lots of rBMP9 were generated in the same cells (Chinese Hamster Ovary, CHO).^15^ Non-reducing SDS-PAGE revealed pro-complex rBMP9 from current studies to be nearly 100% disulfide-linked homodimer at ∼25Kd (lanes 1-2) vs. ∼12.5Kd monomer, with pro-domains migrating at the predicted 40-45 Kd range (**Fig. S2A**), while pro-complex rBMP9 preparations from the original study (lane 3) migrated as ∼25% disulfide-linked homodimers and ∼75% monomers. Under reducing conditions, current (lanes 1-2) and previous preparations (lane 3) all migrated as 12.5 Kd monomers. Despite differential covalent linking of dimers in new versus old rBMP9 preparations, both proteins elicited similar concentration-dependent SMAD1/5/9-mediated gene transcription in microvascular endothelial cells (TIME) in a BMP response element (BRE)-luciferase transcriptional activity assay (**Fig. S2B**). In contrast, a cysteine 329 mutant (rBMP9^C392S^) BMP9 pro-complex protein, expressed as an obligatory monomer due to its inability to form intermolecular disulfide bonds, demonstrated markedly attenuated signaling. To determine how endogenous BMP9 is expressed in the circulation, BMP9 was immunoprecipitated from pooled human AB donor serum using monoclonal anti-BMP9 (MAB3209) to reveal a 25 Kd dimer under non-reducing conditions (“NR”), vs. a single 12.5 Kd species under reducing conditions (“R”) (**Fig. S2C**), supporting the concept that BMP9 is expressed as a disulfide-linked dimer in the circulation. Under non-reducing conditions, preparations of pro-domain complexed, 70% disulfide linked (pro-complex BMP9 WT 70/30), cysteine mutant unlinked (pro-complex BMP9 C329S 0/100), and 100% disulfide linked (pro-complex BMP9 100/0) BMP9 are visualized showing differential abundance of dimer (25 Kd) and monomeric (15 Kd) species in comparison with mature dimeric BMP9 (**Fig. S2D**). All of these species exhibited similar SMAD1/5/9 activation in cultured endothelial cells except the C329S mutant (**Fig. S2E**). While exhibiting similar activity as disulfide-linked homodimer in vitro, we speculate that non-disulfide linked rBMP9 could dissociate into monomers in vivo in various tissue microenvironments, with the potential for competitive inhibition rather than receptor cross-linking and signaling.

### Selective BMP9 inhibition and ACTRIIA-Fc improve hemodynamics and restore pulmonary vascular structure in experimental PH via shared cellular and molecular targets

We compared the impact of anti-BMP9 (MAB3209), ALK1-Fc, and ACTRIIA-Fc (a.k.a., sotatercept)^4^ in the SU-Hx rat model to discern respectively if inhibiting BMP9, BMP9/BMP10, or an array of activin, GDF, and BMP ligands elicits distinct effects. Following the development of severe PH after 3 weeks of SU-Hx, a portion of animals were analyzed after 1 week of treatment under normoxia, while the remainder were analyzed after 3 weeks of treatment (**Fig. 2A**). All three treatments reduced RVSP versus isotype control Ab at 1 week (“Week 4”) and 3 weeks (“Week 6”) of treatment (**Fig. 2B-E**). After 1 week of treatment, ACTRIIA-Fc and anti-BMP9 reduced RV hypertrophy; after 3 weeks of treatment, ACTRIIA-Fc and ALK1-Fc reduced RVH, while anti-BMP9 elicited a trend (p=0.0584) towards improved RVH. All three treatments inhibited pulmonary arteriolar muscularization based on SMA staining (**Fig. 2F-G**). CT pulmonary angiography, revealed restoration of pulmonary vascular density in MAB3209 or ACTRIIA-Fc treated animals as compared to pruning observed with SU-Hx (**Fig. 2H**). Thus, inhibition of BMP9, BMP9/10, and treatment with ACTRIIA-Fc exerted comparable effects on hemodynamics and pulmonary vascular remodeling in this model of PH, and similar to previous observations with ACTRIIA-Fc,^4^ treatment with anti-BMP9 appeared to mitigate loss of vessels in severe angio-obliterative PH.

**Figure 2.**
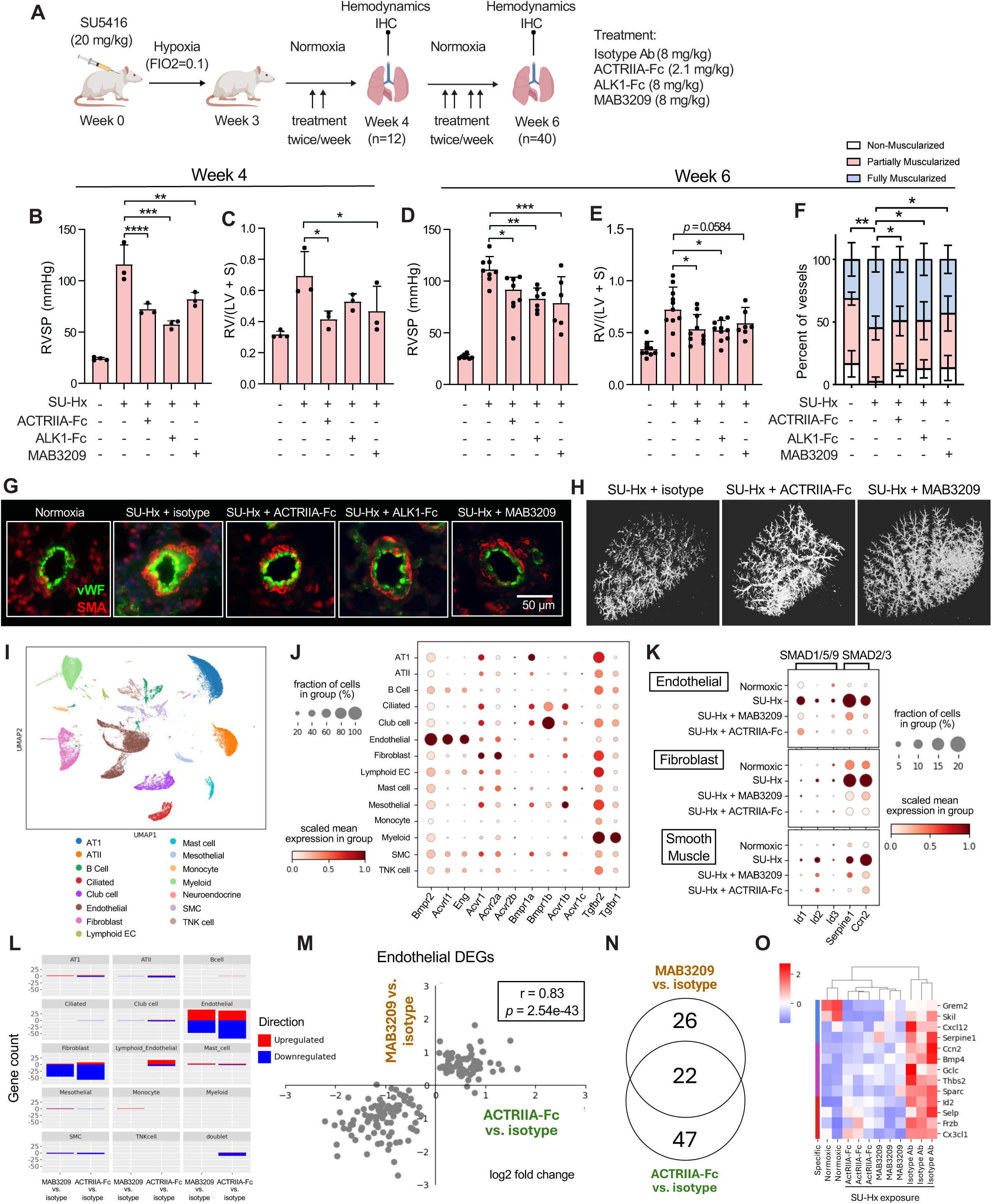
Impact of Anti-BMP9 Ab, ALK1-Fc, and ACTRIIA-Fc on SU-Hx-induced PH in rats. **(A)** Adult male SD rats developed severe PH following exposure to SU-Hx for three weeks, and were treated with anti-BMP9 (MAB3209, 8 mg/kg), ALK1-Fc isotype control (8 mg/kg), ACTRIIA-Fc (2.1 mg/kg), or isotype control Ab (8 mg/kg, all i.p, twice weekly) for 1 or 3 weeks under normoxia. **(B-E)** Treatment with ACTRIIA-Fc, ALK1-Fc, and anti-BMP9 elicited similar improvements in RVSP at 1 week and 3 weeks. ACTRIIA-Fc and anti-BMP9 improved Fulton’s index after 1 week, whereas ACTRIIA-Fc and ALK1-Fc improved Fulton’s index at 3 weeks, and anti-BMP9 elicited a trend (n = 3-11 per group, mean ± SD, *P<0.05, **P<0.01, ***P<0.001, one-way ANOVA with Dunnet’s test (RVSP), and Holm-Sidak’s test (Fulton’s Index). **(F-G)** Immunofluorescence analysis of microvessels (<50 µm) revealed reduced muscularization in SU-Hx rats with anti-BMP9, ALK1-Fc, and ACTRIIA-Fc, shown as mean ± SD, *P<0.05 by one-way ANOVA with Holm-Sidak’s test for the percentage of non-muscularized vessels). **(H)** Micro-CT angiography of the pulmonary vasculature of SU-Hx rats with or without anti-BMP9 treatment reveals improved vascular density with anti-BMP9 treatment. **(I)** Single nucleus RNA-Seq analysis of control and SU-Hx exposed rat lungs reveals broad representations of typical cell populations (doublet population not shown). **(J)** Dot plot of BMP/TGFý receptors across all cell types (doublets and neuroendocrine excluded, expression scaled by column/gene). BMP9/BMP10 receptors Bmpr2, Acvrl1, and Eng are enriched in endothelial lineages; activin receptors Acvr2a and Acvr1b are broadly expressed with enrichment of Acvr2a in fibroblast lineages; TGFβ receptors Tgfbr2 and Tgfbr1 are broadly expressed. **(K)** Dot plot of SMAD1/5/9 and SMAD2/3 target genes in different treatments and cell types (expression scaled by column/gene, within each cell type). At 1 week after return to normoxia, SU-Hx exposure was associated with increased SMAD1/5/9 and SMAD2/3 transcriptional activity based on *Id1/Id2/Id3* and *Serpine1/Ccn2* expression, respectively; One week of treatment with anti-BMP9 or ACTRIIA-Fc elicited similar effects in dampening SMAD1/5/9 and SMAD2/3 transcriptional activity in endothelial and fibroblast lineages. **(L)** Barplots with the number of differentially expressed genes (p_adjust_ < 0.05) for each cluster and contrast. After one week of treatment with anti-BMP9 or ACTRIIA-Fc, the highest number of downregulated genes were in endothelial and fibroblast compartments, whereas the most upregulated genes were found in the endothelial compartment. **(M)** Within endothelium there was a high correlation of genes upregulated and downregulated by anti-BMP9 and ACTRIIA-Fc (r(Spearman)=0.83, p=2.54e-43). **(N)** Within endothelial lineages after 1 week of treatment, the significant differentially expressed genes (DEGs) had a high degree of overlap between anti-BMP9 and ACTRIIA-Fc, with 22 of 48 genes downregulated by anti-BMP9 and 69 genes downregulated by ACTRIIA-Fc shared. **(O)** Heatmap of representative genes (normalized counts, row scaled z-scores) in endothelial cells. Similar samples clusters (columns) are grouped by hierarchical clustering, with representative genes upregulated by exposure to SU-Hx in endothelium and downregulated by treatment with anti-BMP9 and ACTRIIA-Fc shown.

To gain insight into the mechanisms of these treatments, single nucleus RNA-Seq (snRNA-Seq) analysis was performed on tissues after 1 week of treatment to reveal differentially expressed genes (DEG) in various lineages in isotype control treated, anti-BMP9 treated, and ACTRIIA-Fc treated lungs following SU-Hx exposure. UMAP analysis (**Fig. 2I**) revealed distinct clusters corresponding to typical lung cell populations in both control and treated rat lungs. BMP9/BMP10 receptor *Bmpr2* was broadly expressed across lineages, but highly enriched with co-receptors *Acvrl1* and *Eng* in endothelial lineages (**Fig. 2J**). Activin receptors *Acvr2a* and *Acvr1b* were broadly expressed, with enrichment of *Acvr2a* in fibroblast lineages, while TGFβ receptors *Tgfbr2* and *Tgfbr1* were broadly expressed. At 1 week following return to normoxia, SU-Hx exposure was associated with increased SMAD1/5/9 and SMAD2/3 transcriptional activity based on *Id1/Id2/Id3* and *Serpine1/Ccn2* expression, respectively, while one week of treatment with anti-BMP9 or ACTRIIA-Fc dampened SMAD1/5/9 and SMAD2/3 transcriptional activity in endothelial and fibroblast lineages (**Fig. 2K**). Compared to isotype treated control animals, the highest frequency of downregulated genes (**Fig. 2L**) following treatment with anti-BMP9 or ACTRIIA-Fc was observed in endothelial and fibroblast compartments (**Figs. S3, S4**), whereas the most upregulated genes were found in the endothelial compartment, and in the case of ACTRIIA-Fc, the lymphoid endothelial compartment as well. Within endothelium there was a high correlation of log2fold changes of genes upregulated and downregulated by anti-BMP9 and ACTRIIA-Fc (r=0.83, p=2.54e-43, Spearman’s, **Fig. 2M**). Within endothelial lineages, there was a high degree of overlap in downregulated DEGs (p < 0.05, one-sided Fisher’s exact test, **Fig. 2N**) between anti-BMP9 and ACTRIIA-Fc, with 22 of 48 genes downregulated by anti-BMP9 and 69 genes downregulated by ACTRIIA-Fc shared. Representative genes upregulated in endothelium by exposure to SU-Hx and downregulated by treatment with anti-BMP9 or ACTRIIA-Fc are depicted by heat map (**Fig. 2O**) and include *Cxcl12,* encoding stromal derived factor-1 (SDF-1), a predominantly endothelial-derived chemokine known to drive pulmonary vascular remodeling;^20^ *Serpine1,* an early target of SMAD2/3 signaling; *Frzb*, a Wnt decoy receptor and regulator of contractile phenotype in PASMC;^21^ and *Cx3cl1*, a.k.a., fractalkine, an endothelial-derived chemokine and hub gene implicated in PAH-associated inflammation.^22,23^ Taken together these physiologic and molecular data suggests a high degree of overlap between the cellular and molecular mechanisms of anti-BMP9 and ACTRIIA-Fc effects in the rat SU-Hx model, with several shared endothelial-derived targets previously implicated in PAH including *Cxcl12* and *Cx3cl1*.

### High affinity selective anti-BMP9 antibody Ab93 improves experimental PH by attenuating vasoactive factors driving vascular remodeling

A phage-library screening campaign was performed to identify high-affinity human antibodies that competitively inhibit BMP9 signaling by binding to the type II receptor interacting domain. One clone, Ab93, was found to have low picomolar (Kd=54.9 pM) binding affinity for BMP9, based upon surface plasmon resonance (SPR) measurements (**Fig. 3A**). A competitive ELISA to examine the ability of Ab93 to interfere with binding of BMP9 to immobilized type I and type II receptors revealed potent competition with type II receptors ACTRIIA (IC_50_∼0.8 nM), ACTRIIB (IC_50_∼4 nM), and BMPR2 IC_50_∼4 nM), but not type I receptor ALK1 (**Fig. 3B-E**), consistent with binding of Ab93 to the type II receptor domain. Biolayer interferometry confirmed binding of immobilized Ab93 to rBMP9 but none of other ligands tested, including BMP10, activin A, GDF8 and GDF11 (**Fig. S5A**). In contrast, immobilized ACTRIIA-Fc demonstrated binding to BMP9, BMP10, activin A, GDF8 and GDF11 (**Fig. S5B**). Consistent with a high level of selectivity for BMP9 vs. BMP10 in vivo, a sensitive LC-MS method to assay circulating levels of BMP9 and BMP10 in adult male SD rats treated with Ab93 or isotype control antibody (10 mg/kg i.p. twice weekly) revealed ∼5-fold increased levels of circulating BMP9 but no increases in BMP10, consistent with trapping and metabolic protection of circulating BMP9 only (**Fig. S5C-D**). When administered therapeutically in the SU-Hx model for 3 weeks after the establishment of severe PH, Ab93 improved hemodynamics, right ventricular hypertrophy, and pulmonary arteriolar remodeling (**Fig. 3E-H**). To identify potential mechanisms of action for the therapeutic effect of BMP9 inhibition in experimental PH, bulk RNA-sequencing on lung tissues were performed, revealing clustering of treatment (SU-Hx + Ab93) versus disease (SU-Hx + Iso) and control normoxic lung tissues based on principal component analysis (PCA) and depicted by heat maps (**Fig. 3K-L**). In the second principal component axis (PC2), Ab93-treated diseased lung tissues exhibited clustering that was closer to normoxic controls than isotype-treated diseased tissues.

**Figure 3.**
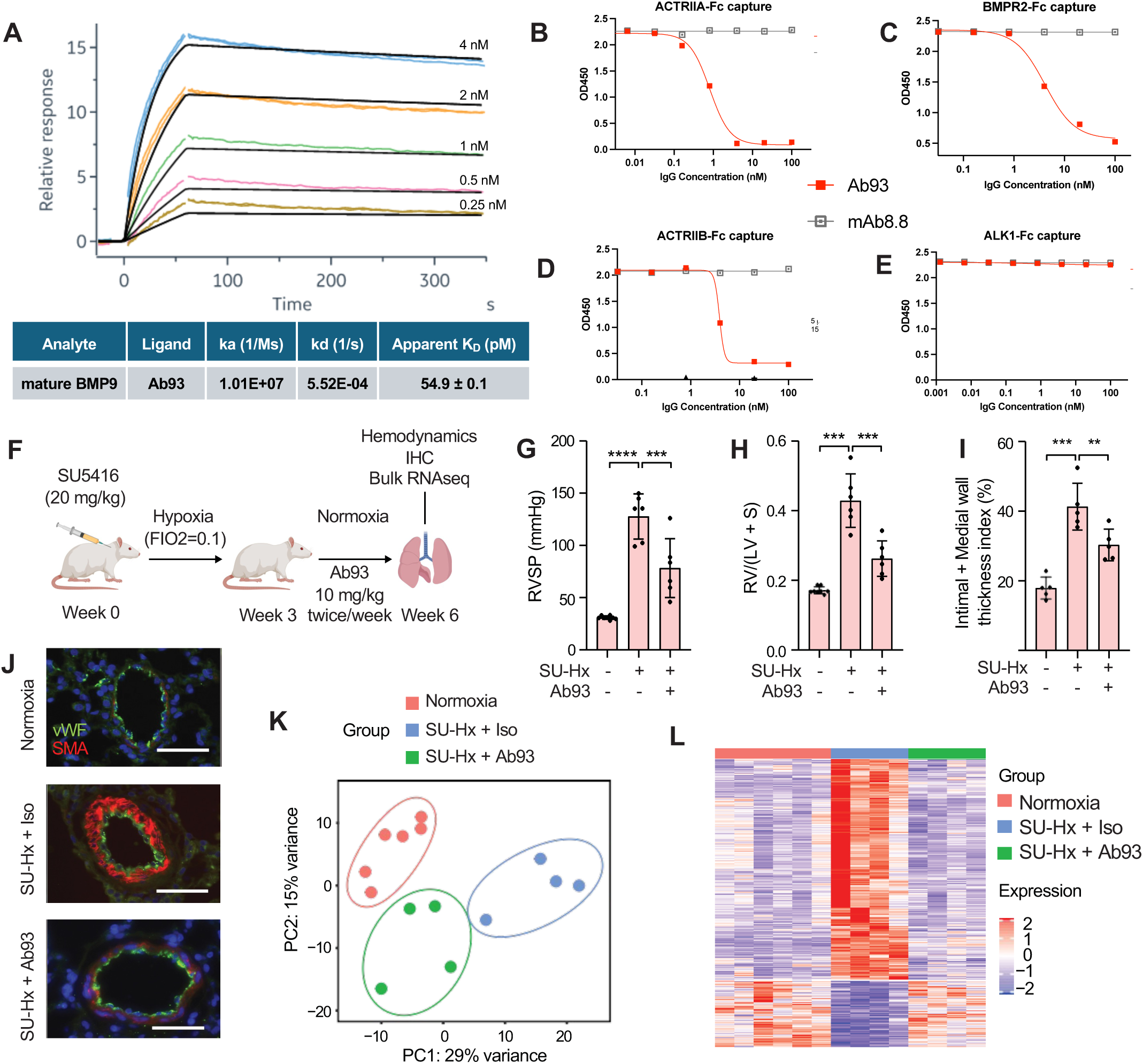
Novel anti-BMP9 antibody Ab93 ameliorates PH in SU-Hx-exposed rats. **(A)** A surface plasmon resonance (SPR) kinetic assay was conducted to measure the affinity of Ab93 to human rBMP9 protein at 37°C via Biacore T200 instrument. Ab93 was adsorbed onto an anti-Fc chip, and rBMP9 was applied with titration series (4 nM to 0.25 nM, with 2-fold dilutions) with 60 s association time and 900 s dissociation time to generate rate constants and affinities, yielding an equilibrium dissociation (K_D_) value of rBMP9 for Ab93 of 54.89 pM. (**B-E**) ELISA measurement of the binding mode of Ab93 for rBMP9. Various type II or type I BMP and activin receptor extracellular domains expressed as IgG Fc fusion proteins (ACTRIIA-Fc, ACTRIIB-Fc, BMPR2-Fc, and ALK1-Fc) were adsorbed onto plates, and incubated with biotinylated rBMP9 (rBMP9-biotin) in the presence of varying concentrations of Ab93 or IgG isotype control (Ab8.8), revealing Ab93 competes with binding of rBMP9 to type II but not type I receptors. (**F**) Adult male SD rats developed severe PH following exposure to SU-Hx for three weeks, and were treated with anti-BMP9 (Ab93, 8 mg/kg) or isotype control (8 mg/kg, both ip, twice weekly) for 3 weeks under normoxia. **(G)** RVSP and **(H)** Fulton’s index were significantly improved in response to treatment (*n* = 6 to 8 per group, mean ± SD, ***P<0.001 by one-way ANOVA with Tukey’s test). **(I-J)** Immunofluorescence analysis of muscularization based on intimal-medial thickness of pulmonary small vessels (< 50 µm) revealed improved muscularization in Ab93 treatment vs. isotype (n = 5 animals per group with 20-30 vessels counted, mean ± SD, **P<0.01, ***P<0.001 by one-way ANOVA with Dunnett’s test). **(K)** Principal component analysis (PCA) of bulk RNAseq analysis of whole lung tissues demonstrated distinct clustering of tissues from animals exposed to normoxia, SU-Hx treated with isotype control, and SU-Hx treated with Ab93. **(L)** Heatmap showing the dysregulated genes in SU-Hx rats’ lung normalized by Ab93.

### Multi-omics analysis reveals several vasoactive endothelial-derived products regulated by BMP9 in lungs from experimental PH and human PAH

To identify specific mechanisms for the therapeutic effect of anti-BMP9 in experimental PH with relevance to human PAH, a single cell RNA-seq dataset from 3 human PAH lungs and 6 unaffected donor lungs,^24^ a bulk RNA-Seq dataset of BMP9-regulated genes in human PMVEC from 5 PAH and 5 unaffected donor lungs,^25^ a bulk RNA-Seq dataset of lungs from Ab93-treated vs. isotype control Ab-treated SU-Hx exposed rats, and normoxic (Nx) rats (**Fig. 4**) were analyzed and compared. Comparison of these datasets allowed us to identify i) DEGs in multiple cell populations relevant to human PAH; ii) DEGs regulated by BMP9 in the pulmonary microvascular endothelium of diseased vs. healthy lungs; and iii) DEGs regulated by inhibition of BMP9 in the SU-Hx model, respectively. Several distinct EC subpopulations, including general capillary (gCap), arterial, systemic venous, pulmonary venous, and aerocyte populations were detected by single cell RNA-Seq of PAH and unaffected lungs, as shown by UMAP plot (**Fig. 4A**). To highlight disease-relevant DEGs in an EC subset implicated in pulmonary microvascular remodeling,^26–28^ gCap ECs were analyzed to reveal 470 genes upregulated in PAH vs. control lungs (“gCap up”, **Fig. 4B; Table S1**). These genes were compared with the 1,219 upregulated genes in SU-Hx vs. Nx control rat lungs (“SU-Hx up”, **Fig. 4B**, **Table S2**), revealing 24 genes were upregulated in both datasets. Among 796 genes that were downregulated in Ab93-treated SU-Hx rats versus isotype control-treated SU-Hx lungs (“Ab93 dn”, **Fig. 4B**), 545 genes were also upregulated in SU-Hx vs. Nx rat lungs, and 8 were also upregulated in gCap ECs of PAH versus unaffected human lungs (**Table S3**). Seven genes were shared between these 3 datasets, including *COL18A1*, *CXCL12*, *IGFBP4*, and *VWA1*. When the 1,196 genes regulated by stimulation of human PMVEC with BMP9 (40 pM, 90 min) were compared with the 470 genes upregulated in gCap ECs of PAH vs. control lungs, 33 genes were shared, including *BMPR2*, *EDN1*, *ENG*, *ID1*, *IGFBP4*, and *VWA1* (**Fig. 4C**), reflecting strong concordance in the list of candidate genes mediating the effects of BMP9 in experimental PH and human PAH obtained from these datasets. To show the relative changes and significance of changes in these candidate genes in the datasets, volcano plots are shown depicting expression of gCap EC genes upregulated in PAH versus controls (**Fig. 4D**), EC genes upregulated by BMP9 stimulation in vitro (**Fig. 4E**), genes regulated by SU-Hx treatment in rats (**Fig. 4F**), and genes downregulated by Ab93 in SU-Hx (**Fig. 4G**). A similar analysis was performed to identify genes normalized by treatment of SU-Hx rats with BMP9/BMP10 trap ALK1-Fc, showing a set of common genes down- and up-regulated by Ab93 and ALK1-Fc including *Ccl21, Grem1, Igf1, Igbp4*, and *Cxcl12* (**Fig. S5E-H**). Taken together these analyses revealed consistency in the genes regulated by BMP9 and BMP9/BMP10 across several datasets from experimental models and human PAH tissues, and shared between several interventions targeting BMP9, BMP9/10, or a broad array of activin, GDF and BMP ligands.

**Figure 4.**
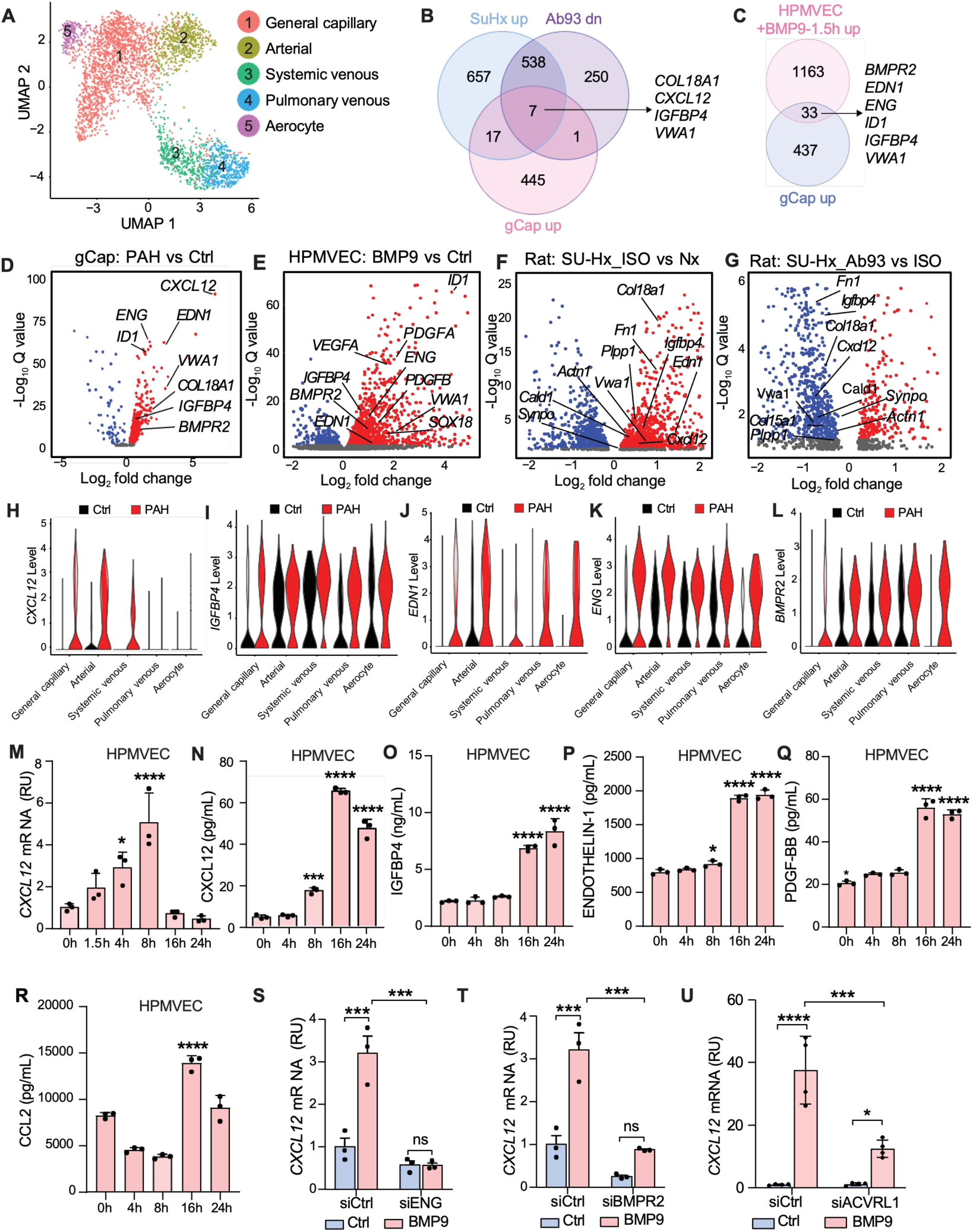
Inhibition of BMP9 attenuates endothelial-derived vasoactive factors driving remodeling in human PAH and experimental PH. **(A)** Uniform manifold approximation and projection (UMAP) plot showing identified endothelial cell types in lungs from 3 IPAH patients and 6 donor controls (GSE169471). **(B)** Venn diagram showing the overlap of upregulated genes in SU-Hx rat lung (SU-Hx–Isotype control vs. Nx), downregulated genes in Ab93-treated SU-Hx rat lung (SU-Hx–Ab93 vs. SU-Hx–Isotype), and upregulated genes in IPAH pulmonary general capillary endothelial cells (gCap) (IPAH vs. Control, GSE169471). **(C)** Venn diagram showing the overlap of upregulated genes in BMP9-treated HPMVEC (BMP9-1.5h vs. control) and upregulated genes in IPAH pulmonary gCap ECs (IPAH vs. Control, GSE169471). **(D-G)** Volcano plots showing changes and significance of candidate driver genes across different datasets, including **(D)** upregulation of *ID1*, *BMPR2*, *CXCL12, EDN1, COL18A1*, and *IGFBP4* in IPAH vs. control lungs; (E) upregulation of *ID1, BMPR2*, *EDN1, IGFBP4, VEGFA, PDGFA, PDGFB, VWA1*, and *SOX18* in BMP9-treated PMVEC; (F) upregulation of *Edn1, Cald1, Vwa1, Col18a1*, and *Igfbp4* in Su-Hx treated rats vs. normoxia; and (G) downregulation of *Cxcl12, Igfbp4, Col18a1, Vwa1*, and *Cald1* in Ab93-treated vs. isotype control treated SU-Hx rats. **(H-L)** Violin plots showing increased expression of *CXCL12*, *IGFBP4*, *EDN1*, *ENG*, and *BMPR2* in various subsets of endothelial cells from IPAH patients (GSE169471). **(M)** BMP9 (40 pM) increased *CXLC12* mRNA expression in HPMVEC in a time-dependent manner. **(N-R)** BMP9 (40 pM) increased secretion of CXCL12, IGFBP4, ET-1, PDGF-BB and CCL2 in cultured HPMVEC supernatants. (*n* = 3 per group, means ± SD. *P<0.05 as compared to 0 h or all the other groups, Dunnett’s test) **(S-U)** BMP9 (40 pM) treatment of HPMVEC increased mRNA expression of *CXCL12*, which was diminished when *BMPR2*, *ACVRL1* or *ENG* were knocked down. (*n* = 3 per group, mean ± SD, \**P*<0.05, \*\**P*<0.01, \*\*\**P*<0.001, \*\*\*\**P*<0.0001 by one-way ANOVA with Sidak’s test).

### *CXCL12*, *IGFBP4*, *EDN1* and *BMPR2* are upregulated by BMP9 in PMVEC, and upregulated in various EC subsets of human PAH

After identifying these candidate BMP9-regulated endothelial genes contributing to experimental PH and human PAH, we analyzed the expression of these transcripts in various EC subsets from IPAH and donor lungs^24^ using consensus markers of 5 lung EC subpopulations^28^ (**Fig. 4H-L; Fig. S6A**). *CXCL12* was upregulated in pulmonary general capillary EC (gCap), as well as in arterial, and systemic-venous EC of PAH vs. control lungs (**Fig. 4H**). *IGFBP4* was expressed in most of these 5 EC subsets, and upregulated in gCap, pulmonary venous, and aerocyte populations of PAH compared to controls (**Fig. 4I**). *EDN1* encoding endothelin-1, a major target of PAH pharmacotherapy, was upregulated in all types of pulmonary ECs of PAH compared to controls, and was particularly abundant in arterial, pulmonary venous and aerocyte populations (**Fig. 4J**). *ENG*, encoding endoglin, a BMP9 co-receptor implicated in heritable PAH and hereditary hemorrhagic telgangiectasia (HHT), and *BMPR2*, the most common locus for heritable PAH, were both upregulated across all clusters of EC in PAH versus controls (**Fig. 4K-L**). *COL18A1*, encoding the precursor protein for endostatin, was upregulated in human PAH and normalized by treatment with Ab93 in SU-Hx rats (**Fig. 4B, D, F, G**). While *COL18A1* was upregulated in arterial endothelial cells in PAH lungs (**S6B-D**), it was not found to be an early transcriptional target of BMP9 in PMVEC (**Fig. 4C, E**) but could be indirectly regulated by BMP9 in the context of disease. Taken together, these multi-omic data implicate several common endothelial-derived, BMP9-regulated vasoactive secreted factors as potential mechanistic drivers of PAH biology.

### *BMPR2*, *ENG* and *ACVRL1* mediate BMP9-mediated CXCL12 expression in a SMAD1/5-dependent manner in PMVEC

Stimulation of cultured PMVEC and PAEC with BMP9 (40 pM) increased levels of *CXCL12* mRNA at 4 and 8 h following treatment, decreasing to basal levels at 16 and 24 h after treatment, and increased accumulation of CXCL12 peptide in culture supernatants at 8 h and peaking at 16 h (**Fig. 4M-N**; **Fig. S6E-F**). Similarly, stimulation of PMVEC or PAEC with BMP9 also increased expression of *IGFBP4* mRNA after 1.5 – 8 h and accumulation of IGFBP4 protein after 16 h (**Fig. 4O; Fig. S3G-H**). Treatment of PMVEC or PAEC with BMP9 increased accumulation of ET-1, PDGF-BB, and CCL2 protein after 16 h (**Fig. 4P-R**; **Fig. S3I-K**). Given the consistency of CXCL12 as an endothelial expressed, BMP9-regulated target across datasets and models, and its known roles in PAH pathobiology and smooth muscle cell plasticity, CXCL12 was explored further as a key mediator of anti-BMP9 therapeutic effect.

Since expression of BMP9 receptors and co-receptors *BMPR2*, *ENG,* and *ACVRL1* are altered in in PAH lungs (**Figs. 4D, 4K, 4L, S7A**) and experimental PAH, we examined their role in BMP9-mediated expression of *CXCL12*. Silencing of *ENG* abrogated the upregulation of *CXCL12* by BMP9 in PMVEC and PAEC (**Figs. 4S**, **S7B**), while silencing of *BMPR2* or *ACVRL1* partially or completely inhibited the expression of *CXCL12* (**Figs. 4T-U**, **S7C-D**). To probe the receptor utilization of BMP9 in regulating *CXCL12* expression further, TIME cells were treated with BMP9 (40 pM) for 3 h in the presence or absence of a panel of siRNA specific for potential BMP9 receptors and co-receptors (**Fig. S7E**). While it was reported that BMP9 can be transduced by *ACVR1*/ALK2 in the absence of *ACVRL1*/ALK1, and by *ACVR2A* in the absence of *BMPR2* in ECs.^29,30^ silencing of *BMPR2*, *ACVRL1*, and *ENG* in TIME cells, but neither *ACVR2A* nor *ACVR1* attenuated transduction of *CXCL12* by BMP9. To ascertain the SMAD effectors responsible for BMP9-mediated expression of *CXCL12*, a panel of siRNA for SMADs 1, 5, 9, 2, and 3 were tested in TIME cells (**Fig. S7F**). Silencing of SMAD1 or SMAD5, but neither SMAD9 nor SMAD2 attenuated BMP9-mediated expression of *CXCL12*. Silencing of SMAD3 resulted in enhanced BMP9-mediated expression of *CXCL12*, suggesting a repressive effect of SMAD3 on *CXCL12* regulation by BMP9.

To examine the potential autocrine or paracrine effects of CXCL12 regulated by BMP9 in ECs, healthy control PMVEC, and PAH donor-derived PMVEC (L104, 256, and 262) were exposed to CXCL12 ligand for 24 h, and the impact on proliferation was assessed by BrdU incorporation. CXCL12 had a potent growth suppressive effect in both donor-derived PMVEC, but this effect was blunted in the PAH derived PMVEC (**Fig. S7G-J**). There was minimal effect of CXCL12 on PAEC proliferation (**Fig. S7K**), while there was a pro-proliferative effect of CXCL12 on PASMC, though less potent than that of PDGF-BB in these cells (**Fig. S7L-M**).

### BMP9-regulated secretion of CXCL12 in PMVEC induces phenotypic modulation in PASMC

To identify potential signaling interactions between PASMCs and PMVECs regulated by BMP9, the overlap in upregulated genes in SU-Hx vs. control lungs that were downregulated by treatment with Ab93, and genes upregulated in PAH lung SMC/pericyte subpopulations was determined (**Fig. 5A, Table S4**). There were 30 genes that were increased in SU-Hx rat lungs, downregulated by Ab93, and upregulated in PAH SMC/pericytes (**Fig. 5B**), including *COL18A1, THY1, FRZB, LTBP2, CPE* and *CNN1–*several of which are markers or regulators of SMC phenotype. Expression of SMC contractile phenotype marker *CNN1* was increased and more frequent in PAH lung-derived SMC/pericytes vs controls (**Fig 5C**). Consistent with the ability of CXCL12 to elicit SMC contractile phenotype in multiple vascular beds and disease contexts.^20,31,32^ stimulation of PASMC with CXCL12 for 24 h elicited expression of contractile phenotype markers *CNN1*, *TAGLN*, *CPE*, *THY1*, and *LTBP2* (**Fig. 5D-H**). Postulating that CXCL12, which was upregulated in PAH-derived gCAP and arterial EC subsets (**Fig. 4H**), we tested whether BMP9 regulates PASMC phenotype directly, or indirectly through the actions of BMP9-CXCL12 signaling in EC. Stimulation of mono-cultured PASMC with BMP9 (40 pM) did not elicit changes in *CNN1* expression after 24 or 48 h (**Fig. 5I**). However, BMP9 stimulation of PMVEC co-cultured on opposite sides of a transwell membrane with PASMC potently elicited *CNN1* expression in PASMC (**Fig. 5J**). This effect of BMP9 on co-cultured PMVEC and PASMC was partly inhibited by co-treatment with anti-CXCL12 antibody, consistent with a paracrine effect mediated by CXCL12 in PMVEC (**Fig. 5K**). Similarly, stimulating co-cultured PMVEC and PASMC with human serum also elicited upregulation of *CNN1* in PASMC (**Fig. 5L**), an effect that was partly abrogated by co-treatment with anti-BMP9 (MAB3209), consistent with inhibition of BMP9 activity in serum. Co-treatment of co-cultured PMVEC and PASMC with CXCR4 inhibitor AMD3100 decreased baseline and BMP9-induced expression of *CNN1* in PASMC (**Fig. 5M**). Integrating data from multi-omic analyses of experimental PH at single cell and single nucleus resolution, human PAH at tissue and single cell resolution, and BMP9 regulated transcription in PMVEC reveals BMP9-mediated regulation of endothelial secreted, vasoactive products including CXCL12, EDN1, IGFBP4, COL18A1, PDGFBB, CCL2, that may contribute to paracine regulation of other vascular cells to drive pulmonary vascular remodeling, serving as targets of anti-BMP9 therapy, and with a high degree of overlap with the mechanisms of ACTRIIA-Fc.

**Figure 5.**
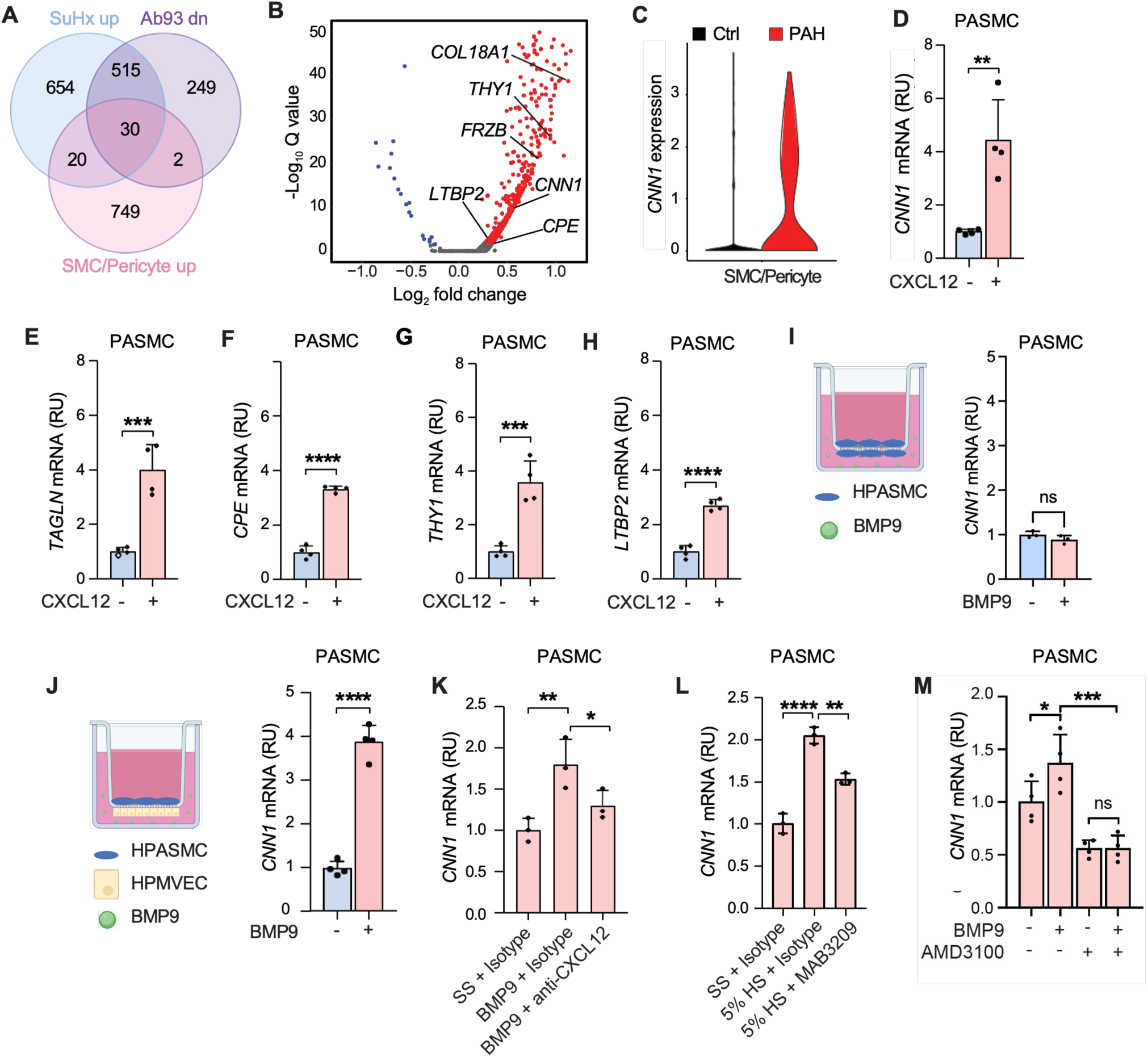
BMP9 stimulates endothelial CXCL12 expression to elicit contractile phenotype markers in co-cultured pulmonary artery smooth muscle cells (PASMC). **(A)** Venn diagram of RNA-Seq analysis of whole lung tissues reveals the overlap of upregulated genes in SU-Hx rat lung vs. normoxic rat lungs (SU-Hx up), downregulated genes in Ab93-vs. isotype control treated SU-Hx rat lung (Ab93 down), and upregulated genes in IPAH pulmonary SMC/pericytes (SMC/Pericyte up, GSE169471). **(B)** Volcano plots depict the expression levels of 6 potentially secreted products of 30 overlapping genes in pulmonary SMC/Pericytes of IPAH patients (GSE169471). **(C)** Violin plot showing significantly increased *Calponin1 (CNN1)* expression in pulmonary SMC/pericyte of IPAH patients (GSE169471). **(D-H)** Treatment with CXCL12 (100 ng/mL) increased expression of contractile phenotype marker genes *CNN1* and *TAGLN*, as well as PAH related genes *CPE*, *THY1*, *LTBP2* in monocultured PASMCs. (*n* = 4 per group, mean ± SD. \*\**P*<0.01, \*\*\**P*<0.001, \*\*\*\**P*<0.0001 by t-test). **(I)** PASMCs did not exhibit increased *CNN1* expression in response to BMP9 (40 pM) when cultured with other PASMC, but when cocultured with PMVEC exhibited increased *CNN1* expression when exposed to BMP9 (*n* = 3 to 4 per group, mean ± SD, \*\**P*<0.01 by t-test. **(J-K)** Treatment of co-cultured PMVEC and PASMC with BMP9 induced *CNN1* expression in PASMC. BMP9 treatment-induced expression of CNN1 in PASMC cocultured with PMVEC **(J)**, which was partially blocked by CXCL12 neutralizing antibody (5 nM, *n* = 3 per group, mean ± SD. \**P*<0.05, \*\**P*<0.01, \*\*\*\**P*<0.0001 by one-way ANOVA with Holm-Sidak test). **(L)** Treatment of co-cultured PMVEC and PASMC with 5% human serum induced *CNN1* expression in HPASMCs, which was partially neutralized by treatment with anti-BMP9 (MAB3209, 5 mM, *n* = 3 per group, \*\**P*<0.01, \*\*\*\**P*<0.0001 by one-way ANOVA with Dunnett’s test). **(M)** BMP9 treatment-induced expression of CNN1 in PASMC that were co-cultured with PMVEC was inhibited by treatment with CXCR4 inhibitor AMD3100 (10 µM, *n* = 4 per group, mean ± SD. \*\**P*<0.01, \*\*\**P*<0.001 by one-way ANOVA with Dunnett’s test).

## Discussion

BMP9 is a pleiotropic growth factor cytokine with context-dependent effects in various developmental and disease contexts. LOF mutations in *GDF2* contribute to PAH in approximately 1% of individuals with group 1 PAH across several large cohorts.^12,33–38^ Consistent with LOF being a heritable risk for PAH, diminished circulating BMP9 is observed in individuals with portopulmonary hypertension or hepatopulmonary syndrome, suggesting that acquired loss of BMP9 due to end-stage liver disease may predispose to PAH.^13,14,39–41^ Based on this rationale, recombinant BMP9 was proposed as a strategy for PAH, with protective effects observed in several preclinical models of PH.^15^ In the current study, a well-characterized recombinant pro-complex BMP9 composed of disulfide-linked homodimeric protein did not recapitulate previously reported therapeutic effects, despite testing at multiple doses in several models. Instead, inhibition of BMP9 or BMP9/BMP10 ameliorated experimental PH, confirming a recent reports implicating BMP9 as a pathogenic driver of pulmonary vascular remodeling,^16,42^ with therapeutic effects when applied before, concurrently, or after the development of experimental PH. In seeking a mechanism for these effects, a wide range of BMP9-regulated, endothelial transcriptional programs are found to overlap with transcriptional changes seen in human PAH, and to be corrected by treatment with anti-BMP9, using a commercial or novel high-affinity specific antibody binding the type II receptor binding domain. Treatment with anti-BMP9 elicited similar degrees of hemodynamic rescue, improvement in RV pathology and attenuation of arteriolar remodeling in experimental PH as ACTRIIA-Fc, with an unexpectedly high degree of correlation (r=0.83) in endothelial DEGs between anti-BMP9 and ACTRIIA-Fc treatment (**Fig. 2K-O**). These findings add our understanding of how BMP9 contributes to the development of PAH as a pathogenic factor, and challenges current notions of how ACTRIIA-Fc/sotatercept, the first-in-class “activin signaling inhibitor” for PAH, achieves its clinical effects.

The observation that inhibition of a single BMP/TGFβ ligand, BMP9, replicates many of the therapeutic and transcriptomic effects of ACTRIIA-Fc, which targets activin A, activin B, GDF8, GDF11, BMP9, BMP10, and several others; **Fig. S5A-B**) suggests either an under-appreciated role of neutralizing BMP9 in the mechanism of action of ACTRIIA-Fc, or a previously unrecognized hierarchical relationship between BMP9 and the other ligands neutralized by ACTRIIA-Fc. Consistent with a potential contribution of inhibiting BMP9 in the mechanism of ACTRIIA-Fc, it was previously reported that inhibition of activin A/B by a non-selective antibody, inhibition of GDF8/11 by a non-selective antibody, or combined inhibition of all 4 ligands in a 4 week SU-Hx prophylaxis model elicited additive improvements in hemodynamics and RV^43^ yet did not match efficacy seen with ACTRIIA-Fc in this model.^4^ In the latter study, BMP9, activin A and GDF11 elicited SMAD1/5 and SMAD2/3 activation in PMVEC, while activin A and GDF11 elicited SMAD1/5 and SMAD2/3 activation in PASMC, suggesting recruitment of both signaling pathways by these ligands. Our current observation that anti-BMP9 and ACTRIIA-Fc both dampened SMAD1/5 and SMAD2/3 signaling across endothelial, fibroblast and smooth muscle lineages in treated lungs (**Fig. 2K**) supports the interpretation that BMP9, activin A and GDF11 may have overlapping effects on signaling in these lineages, with BMP9 making a major contribution.

The impact of ACTRIIA-Fc on BMP9 may have implications for its tolerability. ACTRIIA-Fc was noted to elicit telangiectasia, epistaxis, and GI bleeding in a subset of patients with PAH during phase 2, phase 2 open label extension, and phase 3 trials.^5–7^ These adverse events have been attributed to its ability to bind both BMP9 and BMP10, based on the overlap of these effects with those associated with trials of the BMP9/BMP10 ligand trap ALK1-Fc (dalantercept).^44,45^ In the Phase 2 LUMINA-1 study, an activin A neutralizing antibody (garetosmab) elicited epistaxis in 50% of patients receiving treatment vs. 16.7% receiving placebo during the RCT, and 34.6% of patients receiving treatment in the open label extension, by mechanisms that are currently unknown.^46^ In the present studies, using ACTRIIA-Fc, ALK1-Fc, or anti-BMP9 in experimental PH models was not associated with arteriovenous malformations (AVMs) or associated telangiectasias, bleeding or anemia, and both ACTRIIA-Fc and anti-BMP9 rescued pulmonary vascular density without AVMs (**Fig. 2H**). The animal models, genetic backgrounds, or sizes of cohorts used in this study may be insufficiently sensitive or powered to detect potential adverse effects, while more sensitive models to detect arteriovenous malformations would help assess whether selective targeting of BMP9 might afford a greater therapeutic window as compared to non-selective strategies targeting BMP9, BMP10 and/or activin signaling.

Acquired loss of BMP9 is associated with the development of pulmonary hypertension (PoPH) and hepatopulmonary syndrome (HPS) in patients with end-stage liver disease^13,14,39–41^, raising a theoretical concern that inhibition of BMP9 could provoke this condition in similarly susceptible individuals. In recent studies, it appears that the development of PoPH and HPS are associated with the depletion of both BMP9 <u>and</u> BMP10^13,34^, suggesting loss of both ligands might disrupt a partially redundant function of these ligands. A study comparing the impact of BMP9 vs. BMP10 LOF in mice found that loss of BMP10 exerted more potent and unique effects in promoting arteriovenous malformations that could only be rescued by exogenous BMP10 and not BMP9, suggesting a potentially greater role of BMP10 than BMP9 in maintain capillary vascular structure, and suppression of arteriovenous shunting.^47^ To determine whether or not anti-BMP9 would be a viable therapy for PAH or other vascular diseases will require extensive safety testing in both healthy volunteers, and especially in patients with PAH.

The current studies did not replicate the therapeutic effects of rBMP9 demonstrated in a previous study,^15^ despite using pro-complex protein generated in the same facility from the same cell type and protocol. To ensure these results were not due to variations in dosing or bioavailability, a range of doses that were equivalent, 10-fold, or 50-fold vs. previous doses were tested. Current lots of rBMP9 were essentially 100% disulfide-linked homodimer, in contrast to the original lot which were primarily non-disulfide-linked monomers, and did not elicit changes in hemodynamics, RV, arteriolar remodeling, or whole lung gene transcription in two PH models.

An explanation for the lack of impact of exogenous rBMP9 might be abundant levels of endogenous BMP9 already present in the circulation, previously found to range between 100-250 pg/mL in plasma with no differences between healthy and SU-Hx treated rats, using a sensitive immunoassay^14^. Since the EC_50_ of BMP9 for activation of SMAD1/5/9 and gene transcription in endothelial cells is in the range of 10-20 pg/mL, and human serum has saturated this activity at concentrations of <10%,^14^ we propose that large excesses of exogenous rBMP9 may not elicit additional signaling or biological functions beyond saturation levels of BMP9 signaling elicited by normal rat or human serum. Severely reduced signaling activity of an obligatory non-disulfide linked BMP9 mutant (BMP9^C392S^, **Fig. S2B**) suggests that rBMP9 with a substantial fraction of non-disulfide linked BMP9 could exert inhibitory effects, particularly if non-covalently associated monomers were to dissociate in circulation or in tissues under physiologic conditions. In this study we found that all of the BMP9 that could be immunoprecipitated from serum obtained from a diverse pool of AB blood donors migrated at 25 Kd under non-reducing conditions (**Fig. S2C**). Based on immunoassays using anti-BMP9 capture and anti-BMP10 detection antibodies, or vice versa, it was suggested that circulating BMP9 exists in both homodimeric forms and heterodimeric complexes of BMP9 and BMP10.^48^ In the current study, however, treatment of rats with mono-selective anti-BMP9 antibody Ab93 elicited 5-fold increases in BMP9 plasma concentrations, suggesting sequestration of circulating BMP9, but no increases in circulating BMP10 (**Fig. S5C-D**), suggesting that Ab93 does not interact substantially with a putative circulating BMP9-BMP10 heterodimer.

Beyond its well-described roles in endothelial quiescence and vascular homeostasis,^10,49^ the current study demonstrates critical paracrine regulatory functions of BMP9 in regulating endothelial gene products that could affect the phenotype of other cells including PASMC. The ability of BMP9 to regulate *EDN1*/Endothelin-1 has been described and proposed to contribute to therapeutic effects associated with genetic loss of BMP9 or pharmacologic inhibition of BMP9.^16^ In addition to ET-1, the current study found inhibition of BMP9 is associated with the normalization of several other endothelial-derived vasoactive mediators, several of which have been previously implicated in the pathogenesis of human PAH and experimental PH, including CXCL12, IGFBP4, and COL18A1. These vasoactive secreted proteins have been identified as potential biomarkers of PAH severity and risk^50–52^, and in the case of CXCL12, as a potential mechanistic driver of PH in human disease and experimental models,^53–57^ particularly in its role as an endothelial gene product regulating smooth muscle fate.^20^ CXCL12 was identified as a predominantly endothelial-derived upregulated gene in human PAH lungs by scRNA-Seq (**Fig. 4**)^24^ and is shown here to be normalized by anti-BMP9 in SU-Hx rat lungs by snRNA-Seq and bulk RNA-seq (**Figs. 2, 4, S3**). BMP9 also potently induced in PMVEC the production of PDGF-BB, and CCL2 (**Fig. S3I-J**), which may also contribute to the therapeutic effects of anti-BMP9, although suppression of these transcripts was not observed in treated lungs. The potent therapeutic effects of anti-BMP9 in the current study might be attributed to the large number of vasoactive chemokines, growth factors, and cytokines in the endothelial BMP9 secretome and their downstream effects on vascular cell biology and vascular inflammation. Due to the potentially redundant, complementary, and synergistic effects of these cytokines on vascular cells, as well as the chemoattractant properties for a large variety of immune cells that are known to drive pulmonary vascular remodeling, inhibition of BMP9 as a master regulatory cytokine could represent a potent strategy in the treatment of clinical PAH that may overcome limitations of individual cytokine-directed therapies.

In summary, we have provided several novel and plausible mechanisms by which BMP9 appears to contribute to the pathogenesis PAH, based on the impact of anti-BMP9 in experimental PH on the whole lung and endothelial transcriptome at single nucleus resolution, combined with analysis of the BMP9-mediated secretome in PMVECs from healthy and PAH donors, and a single cell analysis of the endothelial transcriptome in human PAH. We have reconciled our current findings with previous findings that showed a protective effect of exogenous rBMP9, showing a fully disulfide linked BMP9 does not appear to achieve therapeutic effects or alter transcriptional activity in lung tissues of diseased rats. Our findings build upon and extend previous studies that identified mediators of BMP9 impact on endothelial and vascular function, demonstrating broad pro-myogenic, pro-fibrotic, and pro-inflammatory properties on adjacent cells and tissues that might be productively blocked. In the hierarchy of diverse activin, BMP, and other cytokine growth factor targets, BMP9 may be an impactful clinical target due to its increasingly appreciated role as a coordinator of pulmonary vascular remodeling and inflammation due to the multiple downstream signaling cascades that it controls.

## Methods and Materials

### Study design

The objective of this study was to investigate the impact of ligand trap ALK1-Fc, or anti-BMP9 (MAB3209, R&D Systems; Ab93, a human phage-library antibody specific for BMP9 generated for this study) and its targeted ligand (BMP9) and BMP9 target genes (*CXCL12*, *VEGFa*) on pulmonary vascular biology and PH. For in vivo studies, the impact of test agent ALK1-Fc on pulmonary pressures (RVSP) in animal models of PH was considered a primary end point, with impact on RVH and pulmonary arteriolar remodeling considered secondary end points. All animals were segregated into treatment and control groups randomly, and all hemodynamic, physiologic, and histologic measurements were performed in a blinded manner with respect to treatment. Three separate operators at three separate sites performed hemodynamic measurements for independent replication. In some experiments, hemodynamic measurements were performed by direct PA cannulation, whereas in other experiments, the RV was cannulated, depending on which of three independent operators were responsible. On the basis of a hypothesized 30% absolute decrease in RV or PA pressures with ALK1-Fc treatment versus vehicle and a percent coefficient of variation (%CV) of 25% based on previous studies, experimental groups of six animals would be 80% powered to detect this difference, whereas treatment groups of nine animals would be 90% powered to detect this difference. Thus, all animal studies were designed to include groups of 6 to 10 animals at the outset. At the conclusion of each animal study, hemodynamic data were not obtained in some cases because of premature mortality or was excluded from analysis because of prespecified low heart rate threshold (<350 beats/min) during catheterization suggesting a moribund state. Fulton’s index (RV/LV+S) was reported for all animals without exception. Histologic data were obtained by sampling formalin-fixed paraffin-embedded sections from the right middle lobe of the lung or frozen sections from the right lower lobe of the lung after perfusion and fixation. A minimum of 30 to 50 small pulmonary arterioles from each animal subject were photographed and scored in a blinded fashion by two independent observers. Histologic measurements were presented as the mean and SD of measurements across all animals in each cohort and reported without exclusion. For in vitro studies, we hypothesized that BMP9 or its targeted genes would affect a given end point by a minimum of 30%, with a %CV of 15%. Experimental groups consisting of three biological replicates would be 80% powered to this difference, whereas four replicates would be 90% powered to detect this difference. Thus, in vitro studies were designed to include groups of 3 to 4 biological replicates at the outset of each experiment. In some experiments that measured data from cells cultured in multiwell plates, high and low measurements were excluded from all experimental conditions to minimize variability due to differential growth of cells in centers versus edges of plate (plate effect). All animal studies were performed under approved protocols with oversight from the Brigham and Women’s Hospital and Massachusetts General Hospital Institutional Animal Care and Use Committees.

### Reagents

For selected experiments, recombinant “pro-complex” BMP9 was expressed as the pro domain non-covalently associated with a mature human BMP9 homodimer in Chinese hamster ovary (CHO, American Type Culture Collection) cells. Recombinant mature human BMP9 homodimer was purchased from R&D systems. Recombinant ALK1-Fc was expressed as a fusion protein with IgG Fc domain (ALK1-Fc) in CHO cells and purified with two rounds of affinity column chromatography. Monoclonal antibody to BMP9 (MAB3209), and biotinylated affinity purified antibodies to BMP9 (BAF3209) were obtained from R&D Systems. Recombinant CXCL12, PDGF-BB, TGFβ1, and anti-CXCL12 were purchased from R&D Systems. AMD3100 (CXCR4 inhibitor) was purchased from Selleck Chemicals.

### Experimental PH models

Adult male Sprague-Dawley rats (6 to 10 weeks old but uniform within a given experiment; 150 to 170 g, 200 to 250 g, or 400 to 450 g, depending on the cohort) were purchased from Charles River Laboratories. Experimental protocols were approved by the Harvard Institutional Animal Care and Use Committee. Animals were housed at 24°C in a 12-hour light/12-hour dark cycle where food and water were accessible ad libitum. PH was induced by administration of MCT (Oakwood Chemical; 60 mg/kg, sc) followed by 4 weeks of normoxia; or via administration of vascular endothelial growth factor receptor 1/2 (VEGFR1/2) receptor antagonist SUGEN5416 (APExBIO; 20 mg/kg, sc) followed by 3 weeks of normobaric hypoxia (FiO2 = 0.10) and followed by 3 weeks of normoxia.

### Invasive hemodynamic measurements and tissue isolation

Rats were anesthetized with isoflurane (2.5% induction, 0.5 to 1.5% maintenance), and RV pressures were measured by a minimally invasive closed chest approach using a curved tip 2 French pressure transducer catheter (Millar, SPR-513) inserted into the RV through the right internal jugular vein. To assess the degree of RVH, the RV free wall was dissected from the left ventricle plus septum (LV + S) in explanted hearts and weighed. The degree of RVH was determined from the ratio RV/ (LV + S). Lungs were perfused with phosphate-buffered saline (PBS) via the RV to exclude circulating blood cells from the pulmonary circulation, and the right lower lung lobe was ligated, resected, and snap-frozen en bloc for analyses of RNA and protein expression. The pulmonary vasculature was perfused in situ via the RV with 1% paraformaldehyde (PFA) in PBS at 100 cm H2O pressure for 1 min, followed by perfusion of the airways via the trachea at 20 cm H2O for 1 min.

### Pulmonary angiography

In some studies, the right ventricle was canulated with an 18 ga angiocath, advanced to the pulmonary artery position, and perfused with PBS to clear pulmonary vessels, followed by a 50% (v/v) solution of green tissue marking dye (Cancer Diagnostics, Inc.) to opacify the pulmonary arteries. Lungs were then subjected to microCT angiography using a Bruker SkyScan 1176 instrument with a 0.5 mm aluminum filter, 50 kV and 250 µA energy, 295ms exposure time, 18 µm voxel size, and then reconstructed and analyzed using Nrecon and CT analyzer (CTan) software software, with smoothing 4, ring artifact reduction 14, and beam hardening correction of 40% used.

### Lung histomorphometry and immunohistochemistry

To determine the degree of pulmonary vascular remodeling, lungs were embedded for paraffin sectioning or optimal cutting temperature compound sectioning (8 µm) and slides were stained with μ-SMA and von Willebrand factor as previous (11) and visualized by immunofluorescence using an Olympus BX61 epifluorescence microscope. Muscularization of distal intra-acinar vessels (diameter, 10 to 50 µm; 20 to 30 vessels per sample) was scored in a blinded fashion and categorized as either nonmuscular, partially muscularized, or fully muscularized, and relative proportions were expressed as a percentage of total vessels. For fully muscularized intra-acinar vessels, medial wall thickness was calculated on the basis of the following formula: medial thickness − (external diameter − internal diameter)/external diameter × 100.

### Single-nucleus RNA sequencing (snRNA-Seq) sample and library preparation

Frozen rat lungs were pulverized and aliquoted (∼20-50 mg) into 10x Nuclei Isolation 1.5ml tubes and kept on dry ice throughout. Nuclei from each 20-50 mg sample was isolated using the 10x Chromium Nuclei Isolation Kit (10x Genomics, cat #: PN-1000493). Samples were fixed with the plate-based, Parse Biosciences High-Throughput Evercode Nuclei Fixation v3 (part #: ECFN3501) in a 96 well Axygen 450ul V-bottom plate. Samples were processed using the Evercode WT Mega V3 library kit (part #: ECWT3500). Libraries were QC sequenced in-house using the Nextseq 2000. Upon confirmation of quality sequencing data, all libraries were sent to Single Cell Discoveries (SCD) for deep sequencing (25B flow cell) and data was merged for analysis.

### snRNA-Seq data analysis

scRNA-seq dataset (GSE169471, droplet-based data) was re-analyzed by the Seurat R package (4.0.5) in our study^24^. Cells were first filtered with the following criteria: gene number > 200 and < 4,000, and mitochondrial gene percentage < 10. After filtering, a filtered gene-barcode matrix of all samples was integrated to remove batch effects across different donors. In particular, 2,000 shared highly variable genes were identified using Seurat’s ‘SelectIntegrationFeatures()’ function. Integration anchors were identified based on these genes using canonical correlation analysis with 50 dimensions by the ‘FindIntegrationAnchors()’ function. The data were then integrated using ‘IntegrateData()’ and scaled using ‘ScaleData()’. Principal component analysis (PCA) and uniform manifold approximation and projection (UMAP) dimension reduction with 30 principal components were performed. A nearest-neighbor graph using the 30 dimensions of the PCA reduction was calculated using ‘FindNeighbors()’, followed by clustering using ‘FindClusters()’ with a resolution of 0.8. Conserved markers for each cluster were identified using ‘FindAllMarkers()’. Cells were annotated according to the markers identified in original publication^24^. To visualize *CXCL12*, *CNN1* expression among different cell types in the lung, ‘FeaturePlot()’ and ‘VlnPlot()’ were used. For subcluster of endothelial cells, endothelial cell (EC) were subset from the dataset, PCA and UMAP dimension reduction with 50 principal components were performed. A nearest-neighbor graph using the 50 dimensions of the PCA reduction was calculated using ‘FindNeighbors()’, followed by clustering using ‘FindClusters()’ with a resolution of 0.5. Conserved markers for each cluster were identified using ‘FindAllMarkers()’. Cells were annotated according to the markers identified in publications^28,58^. To visualize *CXCL12*, *BMPR2*, *ENG* expression among different endothelial cells in the lung, ‘VlnPlot()’ was used.

The single-cell RNA sequencing data were processed and analyzed using Scanpy (v1.10.3) in Python. Raw count matrices were imported, and cells were filtered based on quality control metrics, including minimum and maximum counts (1,000–10,000), minimum and maximum detected genes (500–3,000), and mitochondrial content (<1%). Genes expressed in fewer than 20 cells were excluded. The data were normalized to 10,000 counts per cell and log-transformed. Highly variable genes (n=4,000) were identified for downstream analysis. Principal component analysis (PCA) was performed, and the top 25 components explaining the majority of variance (>90%) were selected. UMAP was used for dimensionality reduction, and Leiden clustering was applied to identify cell populations. Marker gene overlap and differential expression analysis were conducted to annotate clusters, leveraging literature-based marker genes and external resources such as PanglaoDB and Celltypist.

Differential gene expression analysis was performed using pseudobulk profiles generated by summing raw counts for each cell type and sample. Differential expression analysis was conducted using the DESeq2 framework, implemented via the PyDESeq2 library. Comparisons were made between treatment groups, including ActRIIA-Fc, MAB3209, and isotype controls, across all cell types. Log2 fold changes were calculated, and Wald tests were used to identify significantly differentially expressed genes (adjusted p-value < 0.05).

### RNA isolation, RT-PCR gene expression, bulk RNA Sequencing, and analysis

RNA was extracted from cells or homogenized frozen rat lung samples using TRIzol reagent (15596026, Invitrogen) and dissolved in RNase-free H_2_O. 1 µg total RNA was used to synthesize cDNAs using PrimeScript RT reagents with a genomic DNA elimination reaction (RR047A, Takara). Quantitative (q)PCR was performed using 2.5% of the cDNA product, oligonucleotide primers (Sequences are provided in **Table S5**), SYBR Green Master Mix (A25776, Applied Biosystems), and a thermocycler instrument (QuantStudio 3, Applied Biosystems). The relative mRNA expression levels were determined by the ddC_T_ method and normalized to the relative expression of 18S rRNA. For bulk RNA-Sequencing, total RNA from rat lung tissues (Nx, SU-Hx– Isotype control, SU-Hx–Vehicle, SU-Hx–BMP9 low dose, SU-Hx–high dose)—was extracted using RNeasy Plus Mini Kit (74136, QIAGEN) according to the manufacturer’s instructions with the DNase applied; RNA integrity was assessed using RNA Nano 2000C instrument (Thermo Fisher Scientific) in house and the Bioanalyzer 2100 system (Agilent Technologies, USA) in BGI. The library construction using TruSeq Stranded mRNA Sample Preparation Kit (illumina), library QC and clustering were performed in BGI. After cluster generation, the library preparations were sequenced on the Illumina Novaseq platform (100 bp paired-end reads were generated). RNA sequencing data was transferred from BGI in FASTQ format after filtering by BGI. Paired-end raw reads in fastq format were aligned to the *Rattus norvegicus* rn7 reference genome using the Rsubread R package (2.8.2). FeatureCounts was used to count the read numbers mapped to each gene. Differential expression analysis was performed using the DESeq2 R package (1.34.0). Heatmaps were generated using the ComplexHeatmap R package (2.6.2). Venn diagrams were made using the VennDiagram R package (1.6.20). Volcano plots were generated using ggplot2 R package (3.3.3).

### Affinity of anti-BMP9 Ab93 to human BMP9 determined by SPR

Surface plasmon resonance (SPR) kinetic assay was conducted to measure the affinity of Ab93 to human BMP9 protein at 37°C on a Biacore T200 instrument. Ab93 was captured for 30 sec on an anti-human Fc CM5 sensor. A titration series of human BMP9 protein (4 nM to 0.25 nM, with 2-fold dilutions) was applied to captured Ab93, with 60 sec association time and 900 sec dissociation time. Rate constants and affinities were determined by fitting the resulting sensorgram data to a 1:1 model in Biacore Insight Evaluation software. In a separate experiment, affinity of anti-BMP9 antibody BM01 to human BMP9 was evaluated similarly at 37°C on a Biacore 8K instrument. Same titration series of human BMP9 was used. As summarized in **Fig. 3A**, the equilibrium dissociation (K_D_) values of Ab93 for human BMP9 was 54.89 pM. Representative sensorgrams are shown in **Figure X**.

### RNAi targeting

To knockdown the expression of *BMPR2*, *ACVRL1*, *ENG*, and other genes, HPMVECs were seeded into 4-cm^2^ wells at a density of 2.5 × 10^4^ cells/cm^2^ and transfected with 10 pmol siRNAs when 60% confluent using LipoFectamine RNAiMAX reagent and Opti-MEM (Life Technologies) according to the manufacturer’s instructions. siRNAs targeting *BMPR2* (4390824, Ambion) and negative control siRNA (4390843, Ambion) were purchased from Life Technologies; endoribonuclease-prepared siRNA (esiRNA) targeting *ENG* (EHU091861) or targeting *EGFP* as negative control (EHUEGFP) were obtained from Sigma-Aldrich. RT-qPCR or immunoblotting analysis was used to assess the knockdown efficiency. Cells were starved and treated with rhBMP9 (40 mM) at 24 hours after transfection. For RNA analysis, cells were lysed with 1 mL of Trizol reagent (Life Technologies) per well at 1 or 4 hours after treatment. For protein analysis, culture medium was collected for secreted protein assays, and cells were lysed for immunoblotting analysis with 1× sample buffer (NP0008, Life Technologies) with reducing reagent (NP0009, Life Technologies) at 16 hours after treatment.

### Cell culture

Human PMVECs (CC-2527) and PASMCs (CC-2581) from donors without PH were purchased from Lonza. PMVECs were cultured in complete microvascular endothelial growth medium (EGM-2 MV) supplemented with 1% penicillin-streptomycin and 5% fetal calf serum (Lonza). Additional PMVEC and PASMC were also obtained from explanted human lungs with PAH or unrelated conditions at the time of lung transplant under a Mass General Brigham Institutional Review Board approved protocol as previously described.^4^ Human telomerase immortalized microvascular endothelial (TIME) cells were obtained from ATCC (CRL-4025). PASMCs were cultured in complete smooth muscle cell growth medium (SmGM-2) supplemented with 1% penicillin-streptomycin and 5% fetal calf serum (Lonza). Cell cultures were routinely tested for mycoplasma contamination and used only if negative and otherwise maintained as previously described (8). Serum starvation medium is basal medium with 0.1% FBS without other supplements. Cells were maintained in a 5% CO2-containing, 37°C humidified incubator, passaged using EDTA-trypsin before becoming confluent, and the used between passage 3 to 8. For BMP9 target gene expression analysis, PMVECs were seeded into 4-cm^2^ wells at a density of 2.5 × 10^4^ cells/cm^2^ and were starved and treated with rhBMP9 for different hours when 80% confluent. Culture medium was collected for secreted protein assays and cells were lysed for RNA analysis with 1 mL of Trizol reagent (Life Technologies) per well.

### Human PMVEC-PASMC co-culture system

PMVECs were seeded onto the bottom of the 4.67 cm^2^ upside down Transwell inserts (3428, Costar) at a density of 6.4 × 10^4^ cells/cm^2^ in 600 µL of complete medium for PMVECs. After 2 hours, when cells got adherent, the inserts were inverted, with 1.5 mL and 2.6 mL complete medium for PMVECs in top chamber and bottom chamber, respectively. 24 hours later, PASMCs were seeded onto the top of the Transwell inserts at a density of 1.3 × 10^4^ cells/cm^2^, with all the medium changed to co-culture medium (mixture of PMVECs complete medium and PASMCs complete medium at a ratio of 1:1). When PASMCs were about 60% confluent, cells on both sides were starved with 0.3% FBS in mixture of PMVECs basal medium and PASMCs basal medium at 1:1, followed by exposing PMVECs to 40 pM rBMP9 in starvation medium. 48 hours after rBMP9 treatment, collect PASMC on top and perform RNA isolation using QIAGEN RNeasy mini kit (74104) according to manufacturer’s instructions, followed by above-mentioned RT-qPCR. For control, PASMCs were seeded on both sides of the inserts with the same conditions and procedures.

### Secreted protein assay

To test the concentration of CXCL12, ET1, PDGF-BB, and CCL2 in PMVECs culture medium, the Ella Simple Plex system and the cartridges (Human CXCL12, SPCKB-PS-000299; Human Endothelial-1, SPCKC-PS-000265; HumanCCL2 and PDGF-BB, SPCKC-PS-006510; Protein Simple) were used according to the manufacturer’s instructions. To test the concentration of IGFBP-4 in PMVECs culture medium, the human IGFBP-4 DuoSet ELISA kit (DY804, R&D Systems) was used according to the manufacturer’s instructions.

### Biolayer interferometry

The association of various recombinant BMP, activin and GDF ligands for Ab93 or ACTRIIA-Fc was examined by Bio-Layer Interferometry (BLI) using a Sartorius ForteBio OctetRED 384 instrument. Ab93 or ACTRIIA-Fc were adsorbed onto anti-human IgG Fc binding sensors (AHC). Ligands were detected at concentrations of 100 nM, with a loading time of 300 s, a baseline measurement of 120 s, and association times of 300 s, and dissociation time of 180 s at 37C.

## Acknowledgements

This study was performed with grant funding support from the Pfizer Centers for Translational Innovation (PBY) and from the National Institutes of Health (NIH) National Heart, Lung and Blood Institute (R01HL159443 and R01HL131910, PBY), and National Institute of Arthritis and Musculoskeletal and Skin diseases (R01AR057374, PBY). All of the co-authors were involved in the conception, design, interpretation and analysis of data. (YZ, PY, LT, OK, EF, SR, SL, RS, HJB, KH, EMMH, SPB, CH and PBY) helped to draft and revise the manuscript critically, and all authors provided approval of the submitted manuscript. PBY is a co-founder, SAB member, and stockholder for Keros Therapeutics, which develops therapies for cardiovascular, hematologic, and musculoskeletal diseases targeting bone morphogenetic protein and TGF-β signaling pathways. Several authors (YZ, LT, OVK, KET, SAB, XT, SPB, CH and PBY) are co-inventors on a patent application for PAH-specific therapy that has been filed on behalf of Mass General Brigham and Pfizer, Inc. The interests of PBY are reviewed and managed by Mass General Brigham in accordance with their conflict-of-interest policies.

## Supplementary Materials

### Supplemental Figure Captions

**Figure S1.**
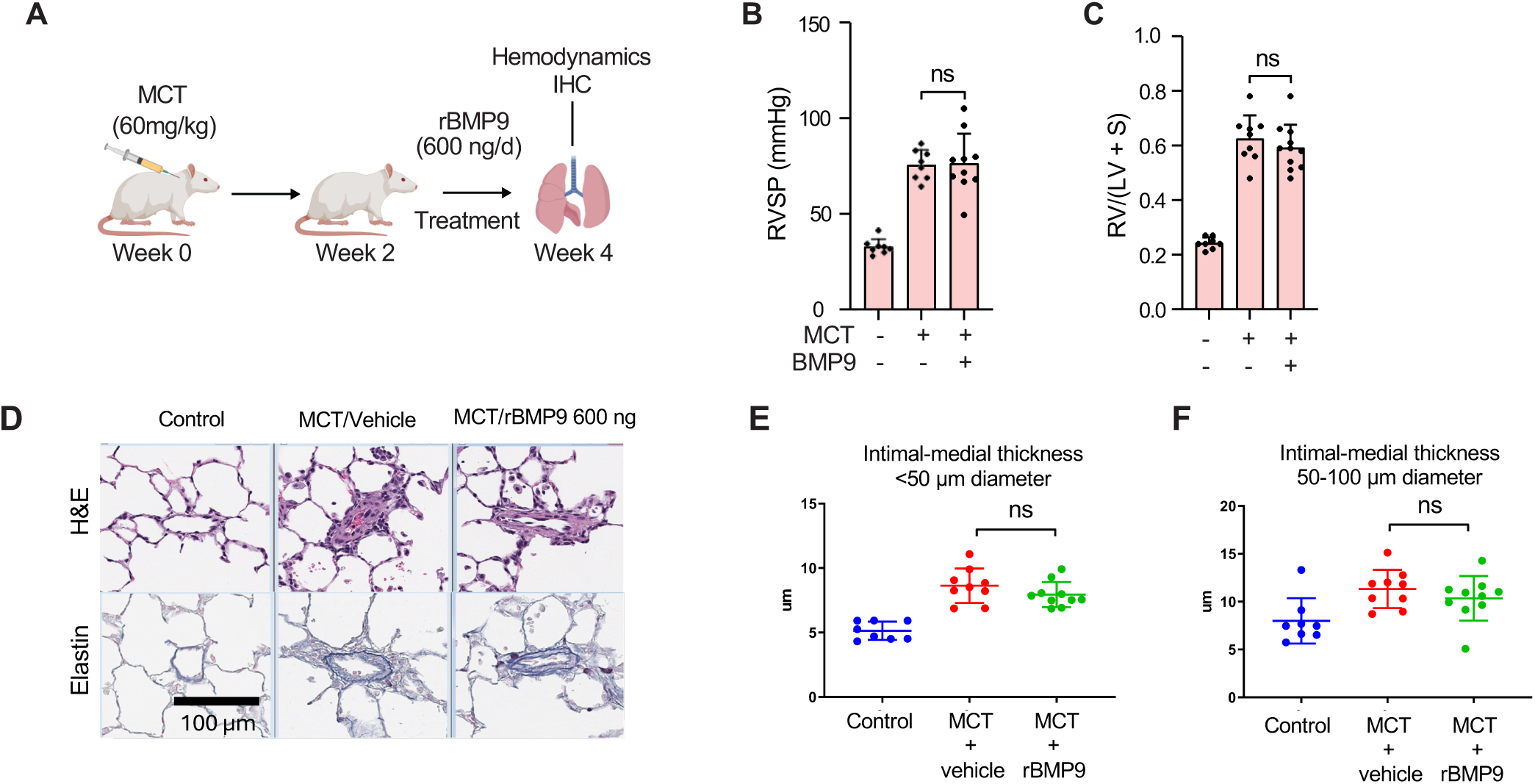
Recombinant pro-complex BMP9 (rBMP9) does not ameliorate PH and vascular remodeling in either MCT or SU-Hx-exposed rats. **(A)** Adult SD rats received MCT (60 mg/kg) and treated with vehicle or rBMP9 (5 µg/kg/d, or 600 ng/d, ip) for 2 weeks starting 2 weeks after MCT, and assessed at 4 weeks for **(B)** RVSP and **(C)** Fulton’s index. **(D)** Elastin histochemical stain revealed MCT-induced vascular remodeling was not mitigated by rBMP9. The vascular intimal-medial thickness of pulmonary arterioles **(E)** <50 µm or **(F)** 50 – 100 µm in diameter was increased in MCT-treated rats and not impacted by rBMP9 vs. vehicle (*n* = 8-11 per group, mean ± SEM, one-way ANOVA with Dunnett’s test).

**Fig. S2.**
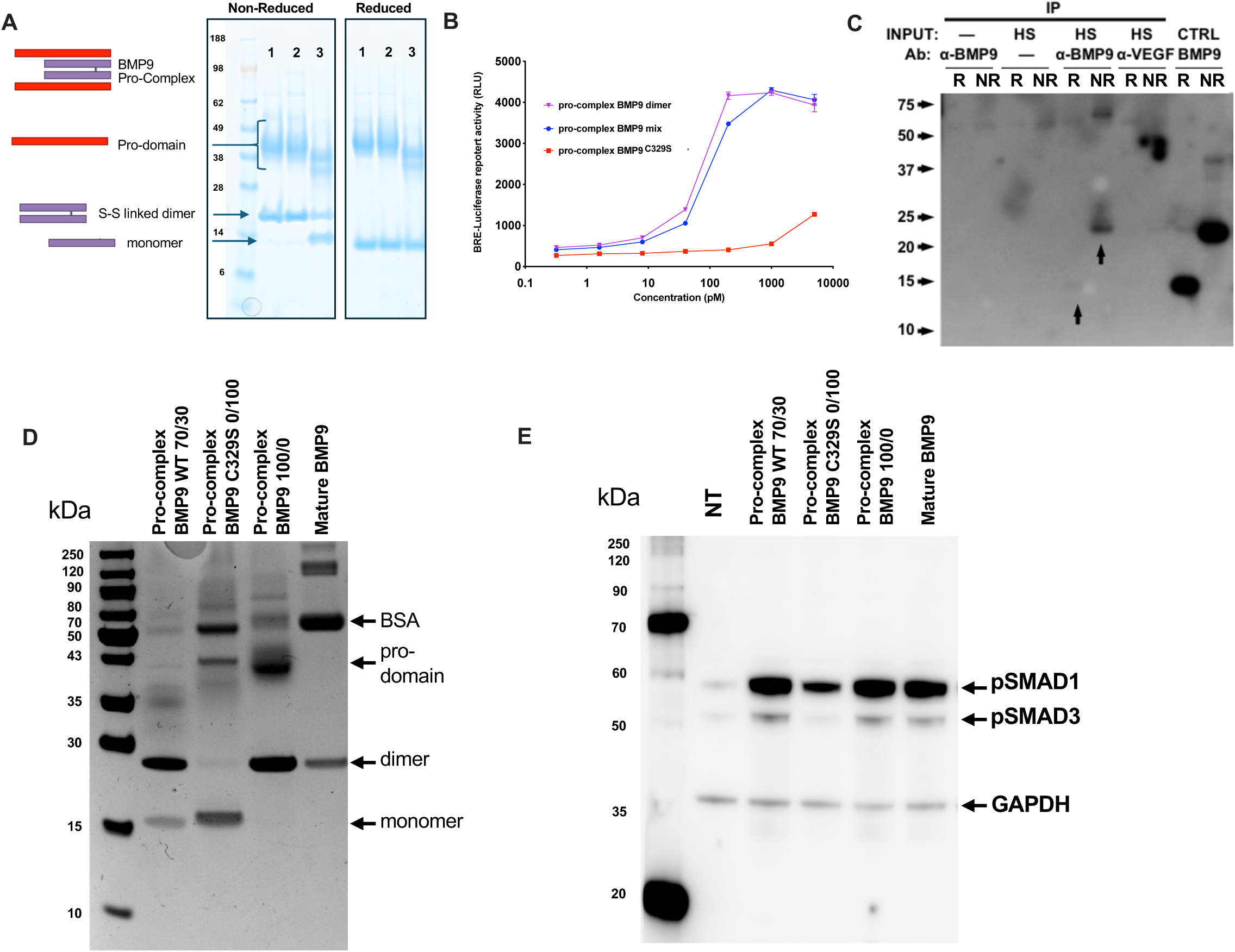
Composition and function of recombinant pro-complex BMP9. **(A)** SDS-PAGE analysis of recombinant pro-complex BMP9 protein from different sources under reducing and non-reducing conditions reveals prodomain of ∼40 Kd, and covalently linked homodimer of ∼25 Kd, and homodimeric from of ∼12.5 Kd. Lanes 1-2 reveal two preparations containing nearly 100% disulfide-linked BMP9 homodimer under non-reducing conditions, both produced in CHO cells; lane 3 reveals ∼25% disulfide-linked homodimer and ∼75% non-disulfide linked BMP9 monomers, also produced in CHO cells and used in reference^15^. **(B)** BRE-luciferase assay showing relative BMP-mediated transcriptional activity in telomerase-immortalized microvascular endothelial cells stimulated for 24 h with 40 pM of 100% disulfide-linked BMP9 dimer, mixed 70%/30% disulfide-linked/non-disulfide linked BMP9 dimer, and 100% non-disulfide linked mutant BMP9^C329S^ protein. **(C)** Immunoblot analysis (biotinylated polyclonal anti-BMP9 BAF3209, followed by streptavidin-HRP) of BMP9 immunoprecipitated from 10 mL pooled human AB donor serum (HS) using anti-BMP9 (MAB3209, α-BMP9, 1 µg/mL, 4C x 12h). Serum immunoprecipitated BMP9 migrated as a ∼25 Kd dimer under non-reducing conditions and as a ∼12.5 Kd monomer under reducing conditions (R), similar to control (CTRL) recombinant mature BMP9 homodimer (1 μg). Anti-VEGFA antibody (α-VEGF) was used as a non-specific immunoprecipitation control antibody. **(D)** SDS-PAGE analysis under non-reducing conditions of recombinant BMP9 produced as 70%/30% disulfide linked/non-linked BMP9 (“70/30”), 100% non-disulfide linked mutant BMP9^C329S^ protein (“0/100”), 100% disulfide linked pro-complex BMP9 (“100/0”), and 100% disulfide-linked mature BMP9. **(E)** Immunoblot demonstrates the ability of different BMP9 preparations to elicit activation of SMAD1 and SMAD3 in cultured TIME cells. A 70%/30% disulfide linked/non-linked BMP9, 100% disulfide linked pro-complex BMP9, and disulfide-linked mature BMP9 elicit similar activation of SMAD1 and SMAD3, whereas non-disulfide linked mutant BMP9^C329S^ (“0/100”) protein elicited attenuated activation of SMAD1.

**Figure S3.**
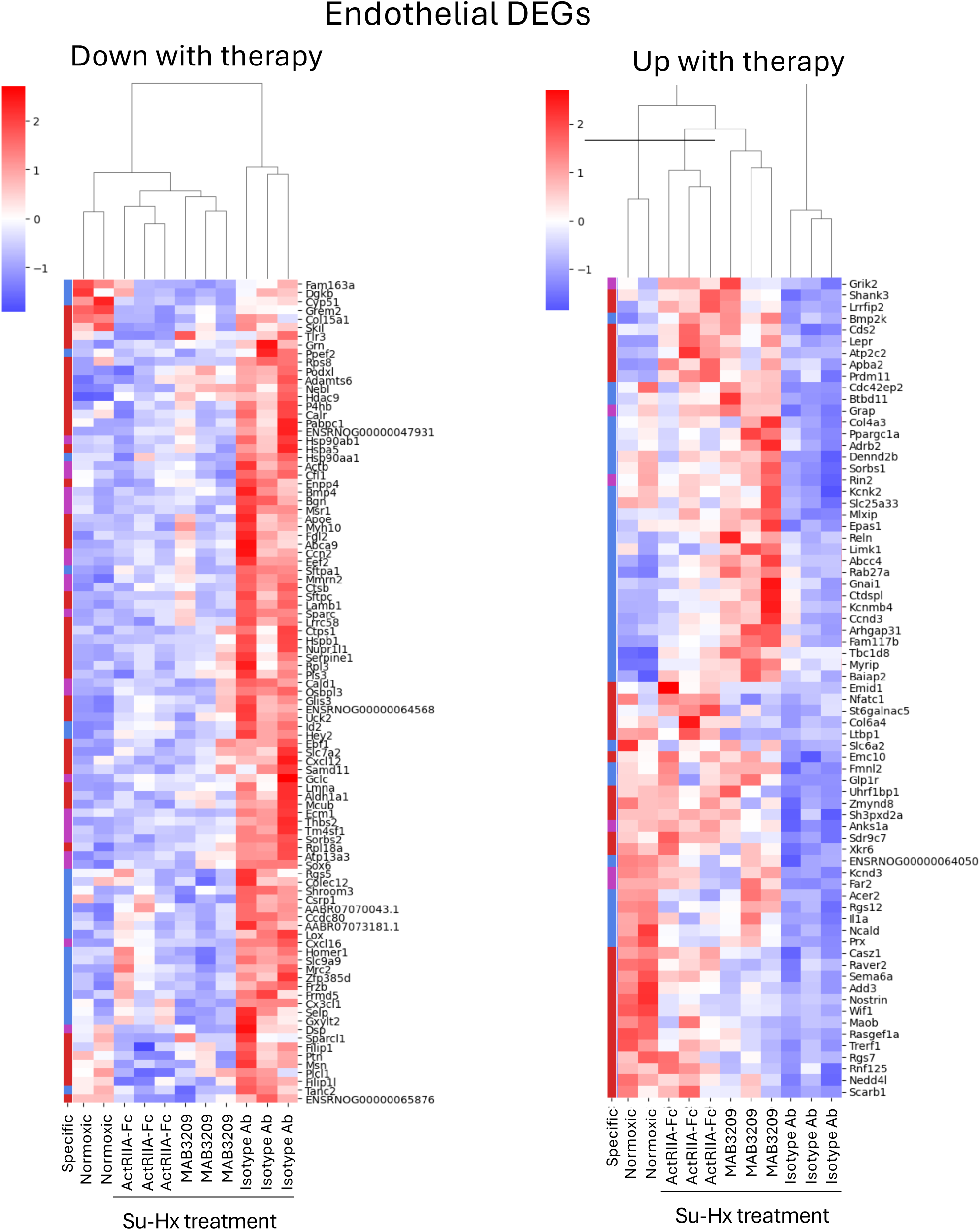
Heatmaps of differentially expressed genes (DEGs) in endothelial cells identified by single nucleus RNA-seq analysis of lung tissues from rats subjected to SU-Hx and various treatments. **(A)** Heatmap of endothelial DEGs between various treatments: anti-BMP9//MAB3209 (8 mg/kg), ACTRIIA-Fc (2.1 mg/kg), or isotype control Ab (8 mg/kg, all i.p, twice weekly) in SU-Hx rats, and normoxic rats. Single-cell expression is aggregated on a sample level, normalized, scaled to z-scores and similar genes (rows) and treatments (columns) aggregated by hierarchical clustering. Row label (‘Specific’) indicates genes that are statistically significant in anti-BMP9 only (red) in ACTRIIA-Fc (blue) only or both treatments (purple). **(A)** Panel denotes endothelial genes downregulated with treatments compared to isotype control. **(B)** Panel denotes endothelial genes upregulated with treatments compared to isotype control.

**Figure S4.**
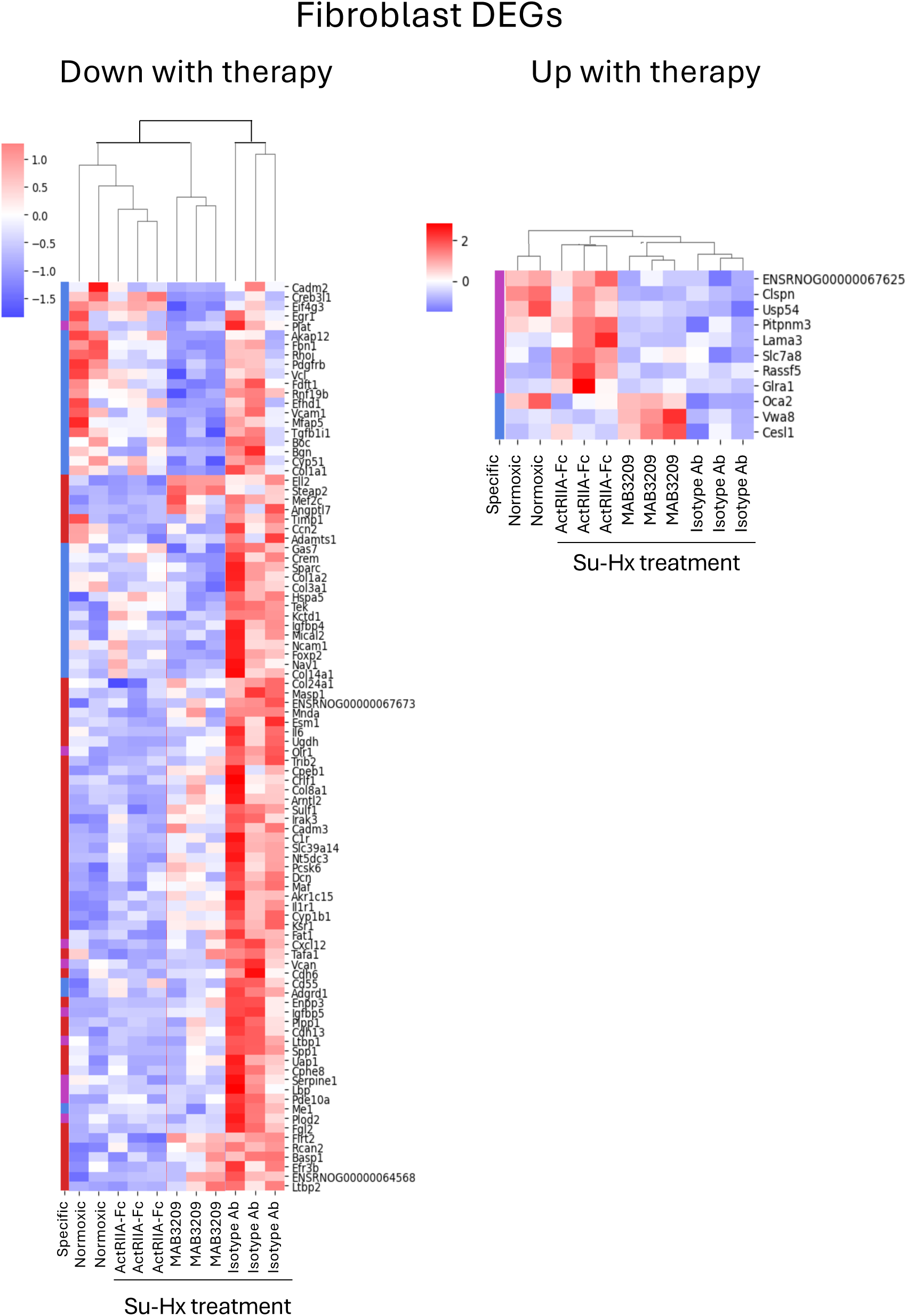
Heatmaps of differentially expressed genes (DEGs) in fibroblasts identified by single nucleus RNA-seq analysis of lung tissues from rats subjected to SU-Hx and various treatments. **(A)** Heatmap of fibroblast DEGs between various treatments: anti-BMP9//MAB3209 (8 mg/kg), ACTRIIA-Fc (2.1 mg/kg), or isotype control Ab (8 mg/kg, all i.p, twice weekly) in SU-Hx rats, and normoxic rats. Single-cell expression is aggregated on a sample level, normalized, scaled to z-scores and similar genes (rows) and treatments (columns) aggregated by hierarchical clustering. Row label (‘Specific’) indicates genes that are statistically significant in anti-BMP9 only (red) in ACTRIIA-Fc (blue) only or both treatments (purple). **(A)** Panel denotes fibroblast genes downregulated with treatments compared to isotype control. **(B)** Panel denotes fibroblast genes upregulated with treatments compared to isotype control.

**Figure S5.**
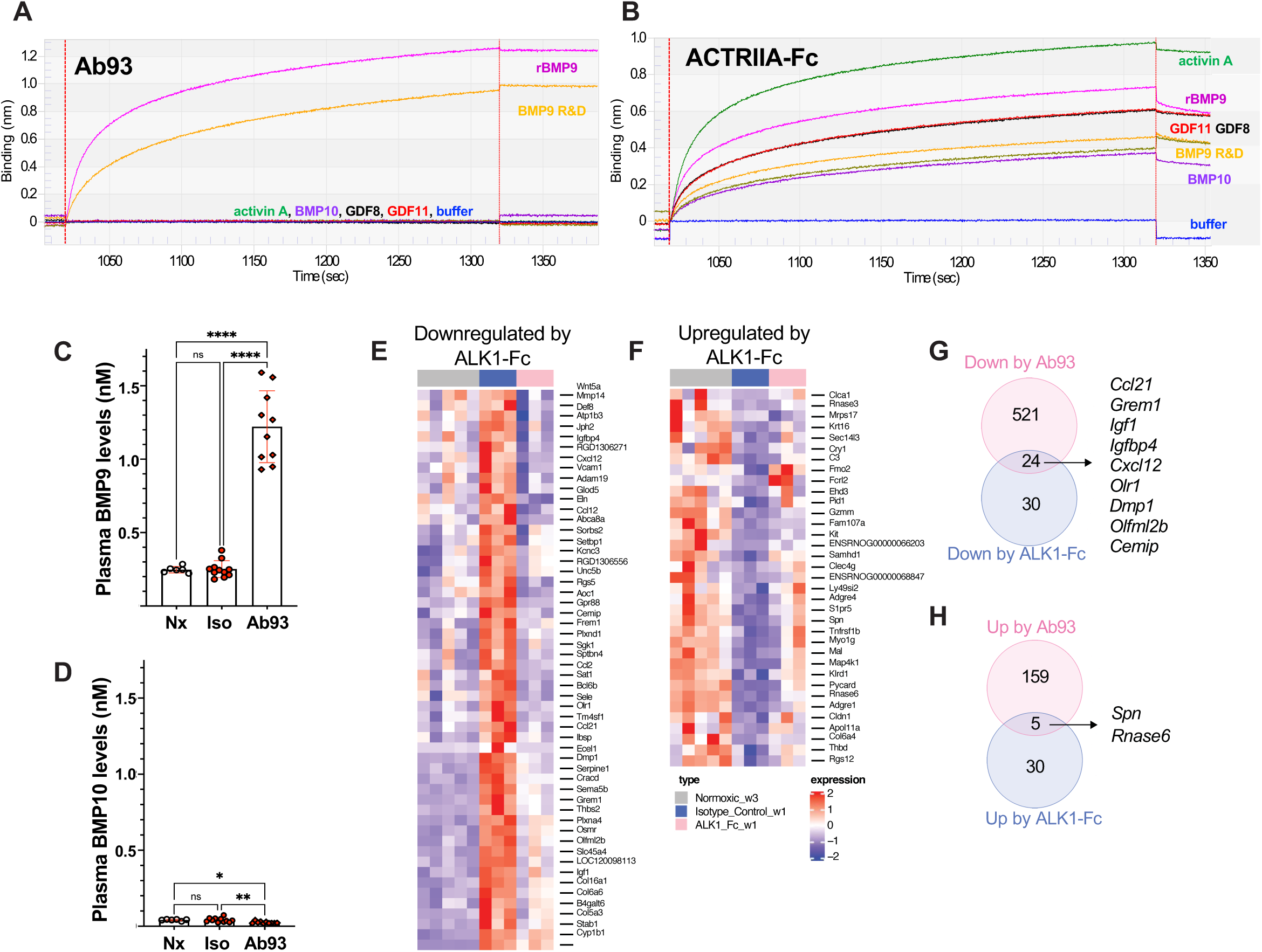
A novel highly selective anti-BMP9 antibody (Ab93) elicits overlapping DEGs in lungs of SU-Hx rats in comparison to treatment with ALK1-Fc. **(A-B)** Biolayer interferometry was used to analyze the ligand specificity of Ab93 and ACTRIIA-Fc. At ligand concentrations of 100 nM, associations of rBMP9 and commercial (R&D) rBMP9 were detected for Ab93 **(A)**, but not other ligands tested (BMP10, activin A, GDF8, GDF11). Under the same conditions, **(B)** interactions for all other ligands tested, rBMP9, BMP10, activin A, GDF8 and GDF11, were found with ACTRIIA-Fc. **(C-D)** Circulating levels of BMP9 and BMP10 were measured by a sensitive LC-MS method in plasma obtained from control adult male SD rats, and those undergoing MCT-induced PH for 3 weeks and treated with Ab93 (or isotype control Ab (10 mg/kg i.p. twice weekly) revealing **(C)** markedly increased levels of circulating BMP9, consistent with trapping and metabolic protection of circulating BMP9, whereas **(D)** levels of BMP10 were not increased but slightly diminished. Values shown are mean ± S.D., *p<0.05, **p<0.01, ***p<0.001, ***p<0.0001, comparisons based on one-way ANOVA with Sidak’s test. **(E-H)** Bulk RNAseq analysis of lungs from SU-Hx exposed rats treated with ALK1-Fc versus Ab93 were compared. **(E)** Heatmap showing DEGs upregulated by SU-Hx in comparison to Normoxic rats and downregulated by ALK1-Fc. **(F) H**eatmap showing DEGs downregulated by SU-Hx in comparison to Normoxic rats and upregulated by ALK1-Fc. **(G)** Comparison of numbers of upregulated DEGs in SU-Hx animals that ere downregulated by Ab93 and ALK1-Fc revealed overlapping sets of genes including *Cxcl12, Igfbp4, Ccl21, and Grem1*. **(H)** Genes that were downregulated in SU-Hx rat lungs that were upregulated by treatment of Ab93 or ALK1-Fc included *Spn* and *Rnase6*.

**Figure S6.**
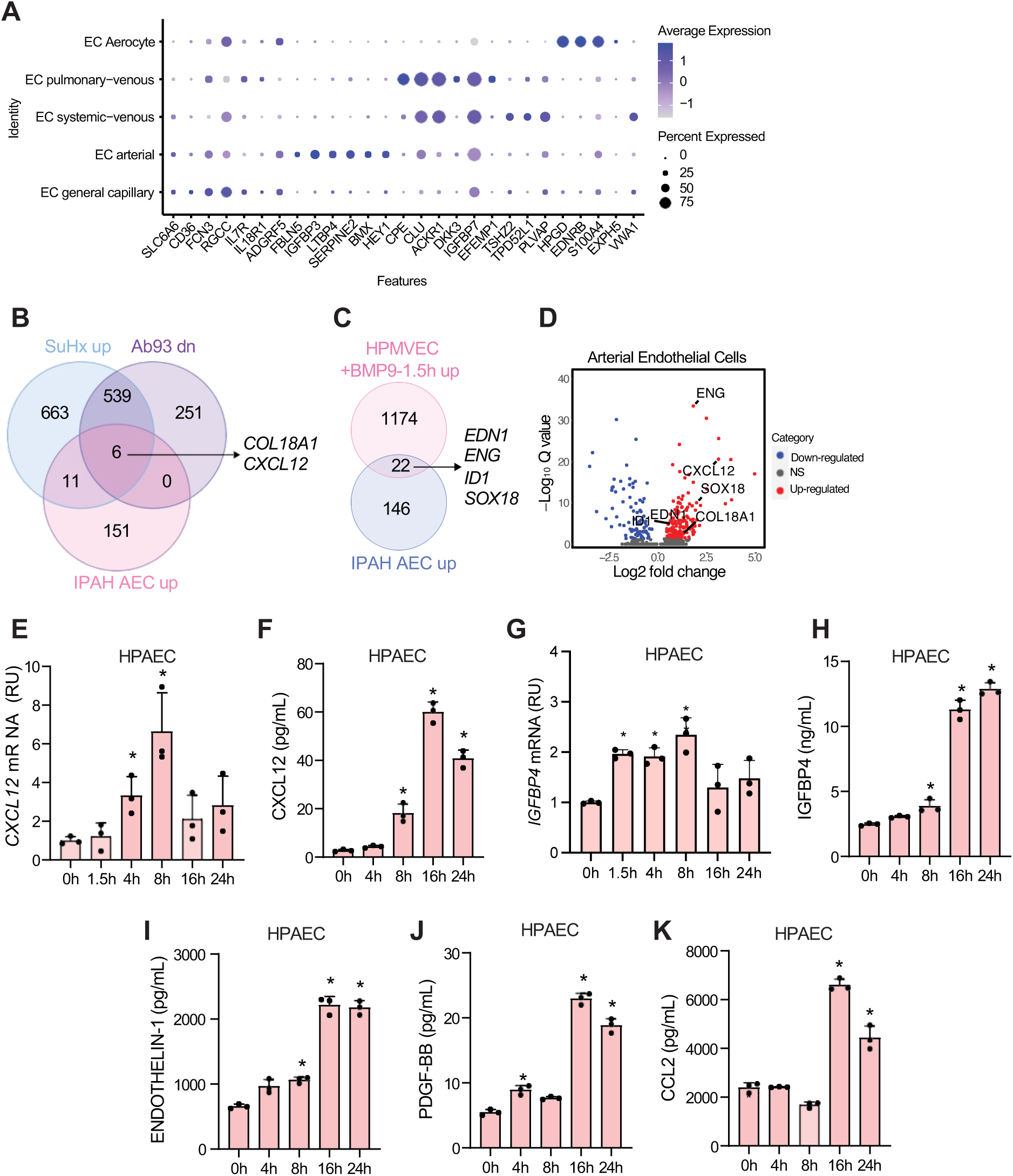
Endothelial subset DEGs in Ab93-treated SU-Hx lung tissues. (**A**) Dot plot highlighting log_10_ average expression of selected marker genes used to identify endothelial clusters. The dot size corresponds to the percentage of cells expressing a gene in a given cluster. (**B**) Venn diagram showing the overlap of upregulated genes in SU-Hx rat lung (SU-Hx–Isotype control vs Nx), downregulated genes in Ab93-treated SU-Hx rat lungs (SU-Hx–Ab93 vs SU-Hx– Isotype), and upregulated genes in IPAH pulmonary artery endothelial cells (AEC, IPAH vs Control, GSE169471). (**C**) Venn diagram showing the overlap of upregulated genes in BMP9-treated HPMVEC (BMP9-1.5h vs control) and upregulated genes in IPAH pulmonary AECs (IPAH vs Control, GSE169471). (**D**) Volcano plot showing potential candidate genes upregulated in IPAH pulmonary AEC (IPAH vs Control, GSE169471). BMP9 (40 pM) increased *CXLC12* mRNA (**E**) and (**F**) protein expression in HPAEC at varying intervals up to 24h. Stimulation of cultured HPAEC with BMP9 (40 pM) increased (**G**) IGFBP4 mRNA and (**H**) protein expression, as well as protein expression of (**I**) Endothelin-1, (**J**) PDGF-BB, and (**K**) CCL2 protein in supernatants at varying intervals over 24h. (*n* = 3 per group, mean ± SD, *P<0.05 as compared to 0 h, one-way ANOVA with Dunnett’s multiple comparisons test).

**Figure S7.**
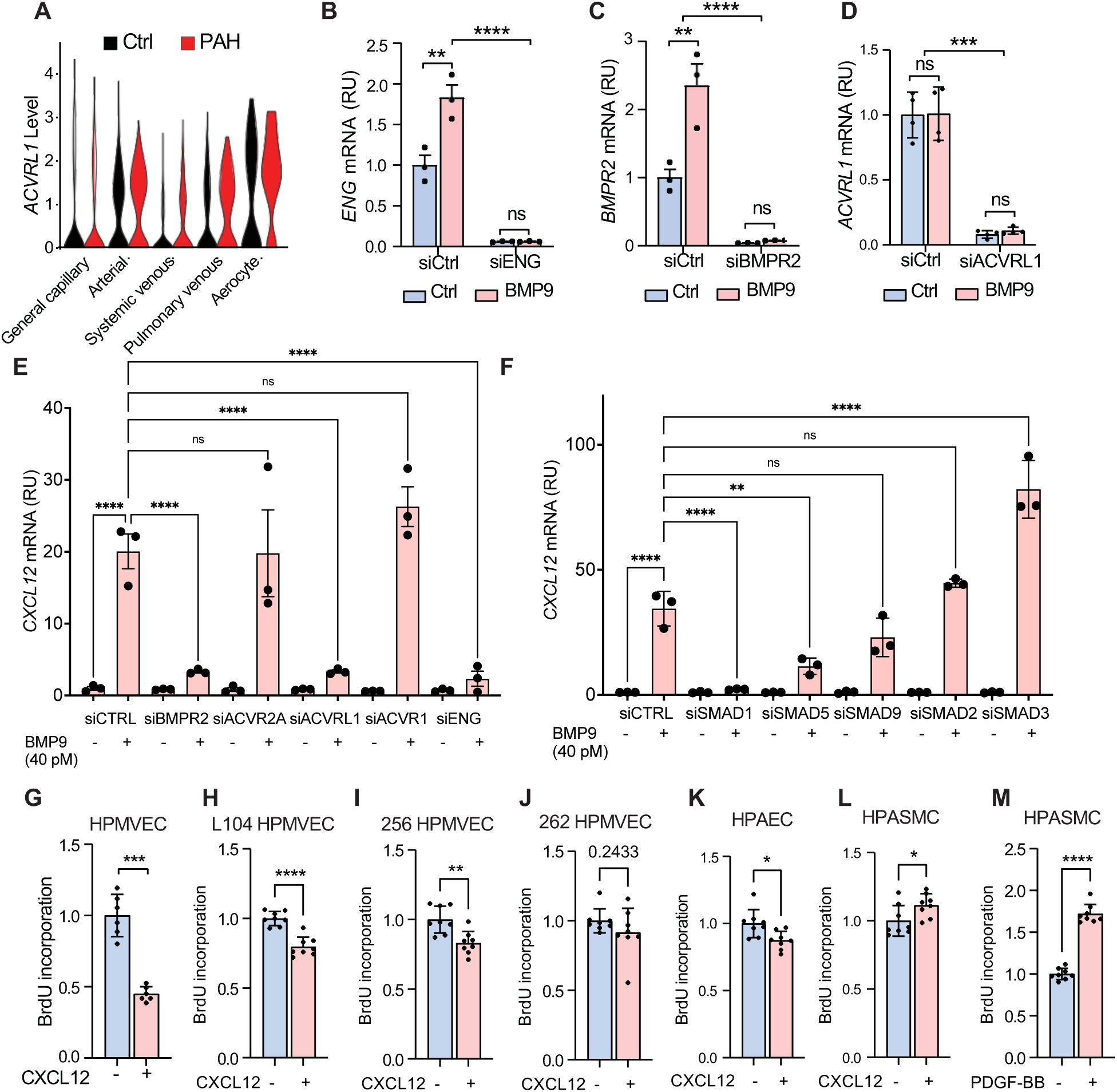
BMP9 regulation of CXCL12 in endothelial cells and impact on proliferation. **(A)** Violin plot showing expression of *ACVRL1* in various subpopulations of human pulmonary endothelial cells (GSE169471). **(B-D)** BMP9 (40 pM) treatment of HPMVEC increased mRNA expression of *ENG* and *BMPR2*, but not *ACVRL1*, each of which were diminished by treatment with specific siRNA (*n* = 3 per group, mean ± SD, \*\**P*<0.01, \*\*\**P*<0.001, \*\*\*\**P*<0.0001 by one-way ANOVA with Sidak’s test). **(E)** BMP9 (40 pM) induced expression of *CXCL12* mRNA in TIME cells in a manner requiring expression of *BMPR2*, *ACVRL1*, and *ENG* based on treatment with specific siRNA. **(F)** BMP9 (40 pM) induced expression of *CXCL12* mRNA in TIME cells in a manner requiring expression of *SMAD1* and *SMAD5* based on treatment with specific siRNA. Values are shown as mean ± S.D., \**P*<0.05, \*\**P*<0.01, \*\*\**P*<0.001, \*\*\**P*<0.0001, or as indicated by one-way ANOVA with Sidak’s test. **(G-M)** BrdU incorporation assay showing that CXCL12 inhibited the proliferation of PMVEC from healthy **(G)** and to a lesser degree PAH donors (**H, I, J)** as well as PAEC **(K)**. CXCL12 and PDGF-BB increased the proliferation of HPASMC (**L** and **M**). (*n* = 8 per group). Values are shown as mean ± S.D., *p<0.05, **p<0.01, ***p<0.001, ****p<0.0001, or as indicated by t-test.

## References

1 Humbert, M. et al. 2022 ESC/ERS Guidelines for the diagnosis and treatment of pulmonary hypertension. Eur Respir J (2022). 10.1183/13993003.008792022

2 Benza, R. L. et al. Predicting survival in pulmonary arterial hypertension: insights from the Registry to Evaluate Early and Long-Term Pulmonary Arterial Hypertension Disease Management (REVEAL). Circulation 122, 164–172 (2010). 10.1161/CIRCULATIONAHA.109.898122

3 Farber, H. W. et al. Five-year Outcomes of Patients Enrolled in the Registry to Evaluate Early and Long-term Pulmonary Arterial Hypertension (PAH) Disease Management (REVEAL). Chest (2015). 10.1378/chest.15-0300

4 Yung, L. M. et al. ACTRIIA-Fc rebalances activin/GDF versus BMP signaling in pulmonary hypertension. Sci Transl Med 12 (2020). 10.1126/scitranslmed.aaz5660

5 Humbert, M. et al. Sotatercept for the Treatment of Pulmonary Arterial Hypertension. N Engl J Med 384, 1204–1215 (2021). 10.1056/NEJMoa2024277

6 Hoeper, M. M. et al. Phase 3 Trial of Sotatercept for Treatment of Pulmonary Arterial Hypertension. N Engl J Med 388, 1478–1490 (2023). 10.1056/NEJMoa2213558

7 Humbert, M. et al. Sotatercept for the treatment of pulmonary arterial hypertension: PULSAR open-label extension. Eur Respir J (2022). 10.1183/13993003.01347-2022

8 Humbert, M. et al. Sotatercept in Patients with Pulmonary Arterial Hypertension at High Risk for Death. N Engl J Med 392, 1987–2000 (2025). 10.1056/NEJMoa2415160

9 David, L., Mallet, C., Mazerbourg, S., Feige, J. J. & Bailly, S. Identification of BMP9 and BMP10 as functional activators of the orphan activin receptor-like kinase 1 (ALK1) in endothelial cells. Blood 109, 1953–1961 (2007). 10.1182/blood-2006-07-034124

10 David, L. et al. Bone morphogenetic protein-9 is a circulating vascular quiescence factor. Circ Res 102, 914–922 (2008). CIRCRESAHA.107.165530

11 Lane, K. B. et al. Heterozygous germline mutations in BMPR2, encoding a TGF-beta receptor, cause familial primary pulmonary hypertension. The International PPH Consortium. Nat Genet 26, 81–84 (2000).

12 Zhu, N. et al. Rare variant analysis of 4241 pulmonary arterial hypertension cases from an international consortium implicates FBLN2, PDGFD, and rare de novo variants in PAH. Genome Med 13, 80 (2021). 10.1186/s13073-021-00891-1

13 Owen, N. E. et al. Reduced circulating BMP10 and BMP9 and elevated endoglin are associated with disease severity, decompensation and pulmonary vascular syndromes in patients with cirrhosis. EBioMedicine 56, 102794 (2020). 10.1016/j.ebiom.2020.102794

14 Nikolic, I. et al. Bone Morphogenetic Protein 9 Is a Mechanistic Biomarker of Portopulmonary Hypertension. Am J Respir Crit Care Med 199, 891–902 (2019). 10.1164/rccm.201807-1236OC

15 Long, L. et al. Selective enhancement of endothelial BMPR-II with BMP9 reverses pulmonary arterial hypertension. Nature medicine 21, 777–785 (2015). 10.1038/nm.3877

16 Tu, L. et al. Selective BMP-9 Inhibition Partially Protects Against Experimental Pulmonary Hypertension. Circ Res 124, 846–855 (2019). 10.1161/CIRCRESAHA.118.313356

17 Theilmann, A. L. et al. Endothelial BMPR2 Loss Drives a Proliferative Response to BMP (Bone Morphogenetic Protein) 9 via Prolonged Canonical Signaling. Arterioscler Thromb Vasc Biol 40, 2605–2618 (2020). 10.1161/ATVBAHA.119.313357

18 Atkinson, C. et al. Primary pulmonary hypertension is associated with reduced pulmonary vascular expression of type II bone morphogenetic protein receptor. Circulation 105, 1672–1678 (2002).

19 Long, L. et al. Altered bone morphogenetic protein and transforming growth factor-beta signaling in rat models of pulmonary hypertension: potential for activin receptor-like kinase-5 inhibition in prevention and progression of disease. Circulation 119, 566–576 (2009). CIRCULATIONAHA.108.821504

20 Dai, Z. et al. Endothelial and Smooth Muscle Cell Interaction via FoxM1 Signaling Mediates Vascular Remodeling and Pulmonary Hypertension. Am J Respir Crit Care Med 198, 788–802 (2018). 10.1164/rccm.201709-1835OC

21 Mocci, G. et al. Single-Cell Gene-Regulatory Networks of Advanced Symptomatic Atherosclerosis. Circ Res 134, 1405–1423 (2024). 10.1161/CIRCRESAHA.123.323184

22 Balabanian, K. et al. CX(3)C chemokine fractalkine in pulmonary arterial hypertension. Am J Respir Crit Care Med 165, 1419–1425 (2002). 10.1164/rccm.2106007

23 Hao, S. et al. Essential Genes and MiRNA-mRNA Network Contributing to the Pathogenesis of Idiopathic Pulmonary Arterial Hypertension. Front Cardiovasc Med 8, 627873 (2021). 10.3389/fcvm.2021.627873

24 Saygin, D. et al. Transcriptional profiling of lung cell populations in idiopathic pulmonary arterial hypertension. Pulm Circ 10 (2020). 10.1177/2045894020908782

25 Szulcek, R. et al. Exacerbated inflammatory signaling underlies aberrant response to BMP9 in pulmonary arterial hypertension lung endothelial cells. Angiogenesis 23, 699–714 (2020). 10.1007/s10456-020-09741-x

26. Liu, B., et al. Single-cell and Spatial Transcriptomics Identified Fatty Acid-binding Proteins Controlling Endothelial Glycolytic and Arterial Programming in Pulmonary Hypertension. bioRxiv (2024). 10.1101/2024.02.11.579846

27. James, J., et al. Novel Populations of Lung Capillary Endothelial Cells and Their Functional Significance. Res Sq (2023). 10.21203/rs.3.rs-2887159/v1

28 Schupp, J. C. et al. Integrated Single-Cell Atlas of Endothelial Cells of the Human Lung. Circulation 144, 286–302 (2021). 10.1161/CIRCULATIONAHA.120.052318

29 Tual-Chalot, S. et al. Endothelial depletion of Acvrl1 in mice leads to arteriovenous malformations associated with reduced endoglin expression. PLoS One 9, e98646 (2014). 10.1371/journal.pone.0098646

30 Upton, P. D., Davies, R. J., Trembath, R. C. & Morrell, N. W. Bone morphogenetic protein (BMP) and activin type II receptors balance BMP9 signals mediated by activin receptor-like kinase-1 in human pulmonary artery endothelial cells. J Biol Chem 284, 15794–15804 (2009). M109.002881

31 Yan, Y. et al. SDF-1alpha/CXCR4 Pathway Mediates Hemodynamics-Induced Formation of Intracranial Aneurysm by Modulating the Phenotypic Transformation of Vascular Smooth Muscle Cells. Transl Stroke Res 13, 276–286 (2022). 10.1007/s12975-021-00925-1

32 Mause, S. F., Ritzel, E., Deck, A., Vogt, F. & Liehn, E. A. Engagement of the CXCL12-CXCR4 Axis in the Interaction of Endothelial Progenitor Cell and Smooth Muscle Cell to Promote Phenotype Control and Guard Vascular Homeostasis. Int J Mol Sci 23 (2022). 10.3390/ijms23020867

33 van den Heuvel, L. M., et al. Genetic Evaluation in a Cohort of 126 Dutch Pulmonary Arterial Hypertension Patients. Genes (Basel) 11 (2020). 10.3390/genes11101191

34 Hodgson, J. et al. Characterization of GDF2 Mutations and Levels of BMP9 and BMP10 in Pulmonary Arterial Hypertension. Am J Respir Crit Care Med 201, 575–585 (2020). 10.1164/rccm.201906-1141OC

35 Zhu, N. et al. Novel risk genes and mechanisms implicated by exome sequencing of 2572 individuals with pulmonary arterial hypertension. Genome Med 11, 69 (2019). 10.1186/s13073-019-0685-z

36 Graf, S. et al. Identification of rare sequence variation underlying heritable pulmonary arterial hypertension. Nat Commun 9, 1416 (2018). 10.1038/s41467-018-03672-4

37 Wang, X. J. et al. Germline BMP9 mutation causes idiopathic pulmonary arterial hypertension. Eur Respir J (2018). 10.1183/13993003.01609-2018

38 Wang, G. et al. Novel homozygous BMP9 nonsense mutation causes pulmonary arterial hypertension: a case report. BMC Pulm Med 16, 17 (2016). 10.1186/s12890-016-0183-7

39 Rochon, E. R. et al. BMP 9/10 in Pulmonary Vascular Complications of Liver Disease. Am J Respir Crit Care Med (2020). 10.1164/rccm.201912-2514LE

40 Robert, F. et al. Disrupted BMP-9 Signaling Impairs Pulmonary Vascular Integrity in Hepatopulmonary Syndrome. Am J Respir Crit Care Med (2024). 10.1164/rccm.202307-1289OC

41 Sangam, S. & Yu, P. B. Hepatopulmonary Syndrome, Another Face of Dysregulated BMP9 Signaling. Am J Respir Crit Care Med (2024). 10.1164/rccm.202405-0895ED

42 Bouvard, C. et al. Different cardiovascular and pulmonary phenotypes for single- and double-knock-out mice deficient in BMP9 and BMP10. Cardiovasc Res 118, 1805–1820 (2022). 10.1093/cvr/cvab187

43 Joshi, S. R. et al. Sotatercept analog suppresses inflammation to reverse experimental pulmonary arterial hypertension. Sci Rep 12, 7803 (2022). 10.1038/s41598-022-11435-x

44 Burger, R. A. et al. Phase II evaluation of dalantercept in the treatment of persistent or recurrent epithelial ovarian cancer: An NRG Oncology/Gynecologic Oncology Group study. Gynecol Oncol 150, 466–470 (2018). 10.1016/j.ygyno.2018.06.017

45 Jimeno, A. et al. A phase 2 study of dalantercept, an activin receptor-like kinase-1 ligand trap, in patients with recurrent or metastatic squamous cell carcinoma of the head and neck. Cancer 122, 3641–3649 (2016). 10.1002/cncr.30317

46 Di Rocco, M. et al. Garetosmab in fibrodysplasia ossificans progressiva: a randomized, double-blind, placebo-controlled phase 2 trial. Nat Med 29, 2615–2624 (2023). 10.1038/s41591-023-02561-8

47 Choi, H. et al. BMP10 functions independently from BMP9 for the development of a proper arteriovenous network. Angiogenesis 26, 167–186 (2023). 10.1007/s10456-022-09859-0

48 Tillet, E. et al. A heterodimer formed by bone morphogenetic protein 9 (BMP9) and BMP10 provides most BMP biological activity in plasma. J Biol Chem 293, 10963–10974 (2018). 10.1074/jbc.RA118.002968

49 Choi, E. J. et al. Enhanced responses to angiogenic cues underlie the pathogenesis of hereditary hemorrhagic telangiectasia 2. PLoS One 8, e63138 (2013). 10.1371/journal.pone.0063138

50 Ambade, A. S. et al. Collagen 18A1/Endostatin Expression in the Progression of Right Ventricular Remodeling and Dysfunction in Pulmonary Arterial Hypertension. Am J Respir Cell Mol Biol (2024). 10.1165/rcmb.2024-0039OC

51 Simpson, C. E. et al. COL18A1 genotypic associations with endostatin levels and clinical features in pulmonary arterial hypertension: a quantitative trait association study. ERJ Open Res 8 (2022). 10.1183/23120541.00725-2021

52 Torres, G. et al. Insulin-like growth factor binding Protein-4: A novel indicator of pulmonary arterial hypertension severity and survival. Pulm Circ 13, e12235 (2023). 10.1002/pul2.12235

53 Montani, D. et al. C-kit-positive cells accumulate in remodeled vessels of idiopathic pulmonary arterial hypertension. American Journal of Respiratory & Critical Care Medicine 184, 116–123 (2011).

54 Xu, J. et al. Inhibition of CXCR4 ameliorates hypoxia-induced pulmonary arterial hypertension in rats. Am J Transl Res 13, 1458–1470 (2021).

55 Bordenave, J. et al. Neutralization of CXCL12 attenuates established pulmonary hypertension in rats. Cardiovasc Res 116, 686–697 (2020). 10.1093/cvr/cvz153

56 Yuan, K. et al. Mural Cell SDF1 Signaling Is Associated with the Pathogenesis of Pulmonary Arterial Hypertension. Am J Respir Cell Mol Biol 62, 747–759 (2020). 10.1165/rcmb.2019-0401OC

57 Yi, D. et al. Endothelial Autocrine Signaling through CXCL12/CXCR4/FoxM1 Axis Contributes to Severe Pulmonary Arterial Hypertension. Int J Mol Sci 22 (2021). 10.3390/ijms22063182

58 Gillich, A. et al. Capillary cell-type specialization in the alveolus. Nature 586, 785–789 (2020). 10.1038/s41586-020-2822-7

